# The PRMT5-splicing axis is a critical oncogenic vulnerability that regulates detained intron splicing

**DOI:** 10.1101/2024.12.17.628905

**Authors:** Colin E. Fowler, Natalie A. O’Hearn, Griffin J. Salus, Arundeep Singh, Paul L. Boutz, Jacqueline A. Lees

## Abstract

Protein arginine methyltransferase 5 (PRMT5) is a promising cancer target, yet it’s unclear which PRMT5 roles underlie this vulnerability. Here, we establish that PRMT5 inhibition induces a special class of unspliced introns, called detained introns (DIs). To interrogate the impact of DIs, we depleted CLNS1A, a PRMT5 cofactor that specifically enables Sm protein methylation. We found that many, but not all, cell lines are CLNS1A-dependent and established that loss of viability is linked to loss of Sm protein methylation and DI upregulation. Finally, we discovered that PRMT5-regulated DIs, and the impacted genes, are highly conserved across human, and also mouse, cell lines but display little interspecies conservation. Despite this, human and mouse DIs have convergent impacts on proliferation by affecting essential components of proliferation-regulating complexes. Together, these data argue that the PRMT5-splicing axis, including appropriate DI splicing, underlies cancer’s vulnerability to PRMT5 inhibitors.

## Introduction

PRMT5 is the major type II arginine methyltransferase and the primary enzyme responsible for catalyzing the symmetric dimethyl arginine mark (SDMA)^1^. PRMT5 is highly expressed in stem cells, and its expression decreases as cells enter post-mitotic, terminally differentiated fates^2^. Importantly, PRMT5 is re-expressed in nearly all cancers, and chemical or genetic inhibition shows drastic anti-proliferative effects^1,3^. Many PRMT5 inhibitors (PRMT5i) show promising preclinical results in a wide variety of tumor types^4–7^, and, as such, several PRMT5i are already in, or moving towards, clinical trials^8^.

PRMT5 writes the SDMA mark on proteins involved in cellular processes including transcription, translation, and splicing. PRMT5 functions in a hetero-octameric complex comprised of four PRMT5 and four cofactor molecules^9,10^. One of these cofactors, MEP50, is required for efficient PRMT5 activity. The other co-factors, CLNS1A, RIOK1, and COPR5, augment PRMT5 activity by recruiting the complex to specific targets associated with splicing, translation, or transcription, respectively^1,11^. The PRMT5-MEP50-CLNS1A complex, referred to as the methylosome^12,13^, is the most studied of these complexes. It binds to and promotes the methylation of three Sm proteins: SmB/B’, SmD1, and SmD3^12,14–16^. These methylated Sm proteins are then incorporated into a larger, seven-member Sm protein ring, which is loaded around an snRNA to form an snRNP – a core component of the spliceosome^17,18^. PRMT5 in known to methylate other splicing factors^19^, but does so without binding to CLNS1A.

Which aspect(s) of PRMT5 biology underly the vulnerability of cancer cells to PRMT5i remains a key question in the field. Because the methylosome is PRMT5’s best understood role, much attention has been focused on assessing splicing problems upon PRMT5 inhibition. In a seminal study, Bezzi et al. examined PRMT5 deletion in neural stem and progenitor cells, and showed that this was sufficient to cause neonatal inviability^20^. They showed that the PRMT5- deficient neural cells display an array of splicing defects, highlighting inclusion of a cassette exon in *MDM4* and consequent p53 activation, but also noting elevated levels of unspliced introns^20^. In a different study, we identified *PRMT5* as the top hit in an *in vivo* screen for essential genes in glioblastoma, and our molecular analysis revealed that the predominant impact on splicing was increased levels of introns^2^. Notably, based on their relative stability and presence in poly-adenylated mRNAs, we concluded that these were likely a special class of unspliced introns, called detained introns (DIs)^2^. DIs occur in fully processed transcripts that remain in the nucleus, making them unavailable for translation^21^. This distinguishes them from retained introns (RIs), which exist in transcripts that reach the cytoplasm and typically undergo nonsense-mediated decay (NMD)^21^. It is now common to use splicing defects as evidence of PRMT5i action^2,4–7,19,22–25^, reinforcing the prevailing view that PRMT5’s splicing role largely underlie the vulnerability of cells to PRMT5 inhibition. However, it is important to note that previous studies use genetic and/or pharmacological approaches that disrupt all PRMT5 functions. Therefore, while splicing defects are evident, it is an open question whether these are responsible for PRMT5i sensitivity versus other roles of PRMT5. Intriguingly, a recent study showed that bulk depletion of CLNS1A, the methylosome co-factor, reduced the viability of pancreatic ductal adenocarcinoma KP4 cells^22^. This observation is consistent with a key role for the PRMT5-splicing axis, but, unfortunately, this study did not look at the impact of CLNS1A loss on splicing.

In this current study, we conduct in-depth analyses of PRMT5-regulated splicing events and directly test their essentiality. We demonstrate that elevated intron levels are the predominant splicing defect and confirm that these are *bona fide* DIs. We then probe the functional importance of the methylosome, independent of other PRMT5 functions, through CLNS1A depletion. We find that CLNS1A is essential for viability in some, but not all, cell lines. Importantly, loss of viability directly reflects loss of Sm protein methylation and upregulation of DIs. Finally, we find that the identities of PRMT5-regulated DIs and the affected genes are highly conserved across human cell lines, as well as mouse cell lines, irrespective of tissue/tumor type. In contrast, there is little conservation at the DI or gene level between human and mouse DIs, even within a single tissue. Despite this, human and mouse DI-containing genes strongly converge on biological processes associated with cell proliferation. This reflects the fact that many DI-regulated genes encode for proteins that are essential, but largely non-overlapping, components of multiprotein complexes that play central roles in proliferation. Together, these data suggest that maintaining appropriate DI splicing in core proliferation-associated genes plays a critical role by which PRMT5 enables cell viability.

## Results

### PRMT5-regulated intronic splicing events are *bona fide* DIs

We began our study by analyzing the splicing changes resulting from PRMT5 inhibition.

We conducted a 3-day 10 nM PRMT5i treatment using the SAM-competitive inhibitor, JNJ- 64619178^6^, in the PRMT5i-sensitive U87 glioblastoma cells^2^. We enriched for poly-adenylated mRNAs in whole cell extracts, performed deep sequencing (>40M reads/sample), and conducted analyses to identify and quantify PRMT5-regulated splicing changes. The majority of significant changes were elevated levels of introns, with the remainder being low levels of exon-based splicing events [cassette exons (CAS), mutually exclusive exons (MXE) and alternative 5’ (A5PSS) or 3’ (A3PSS) splice sites] (**Figure 1a-c**, p_adj_<0.05). We wanted to establish that the persisting introns were *bona fide* DIs, not RIs. A defining hallmark of DI-containing transcripts is their nuclear localization^21^. Thus, we repeated the 3-day 10 nM PRMT5i treatment of U87 cells and performed cellular fractionation, confirming efficacy by western blotting (**Figure S1a-b**).

**Figure 1.**
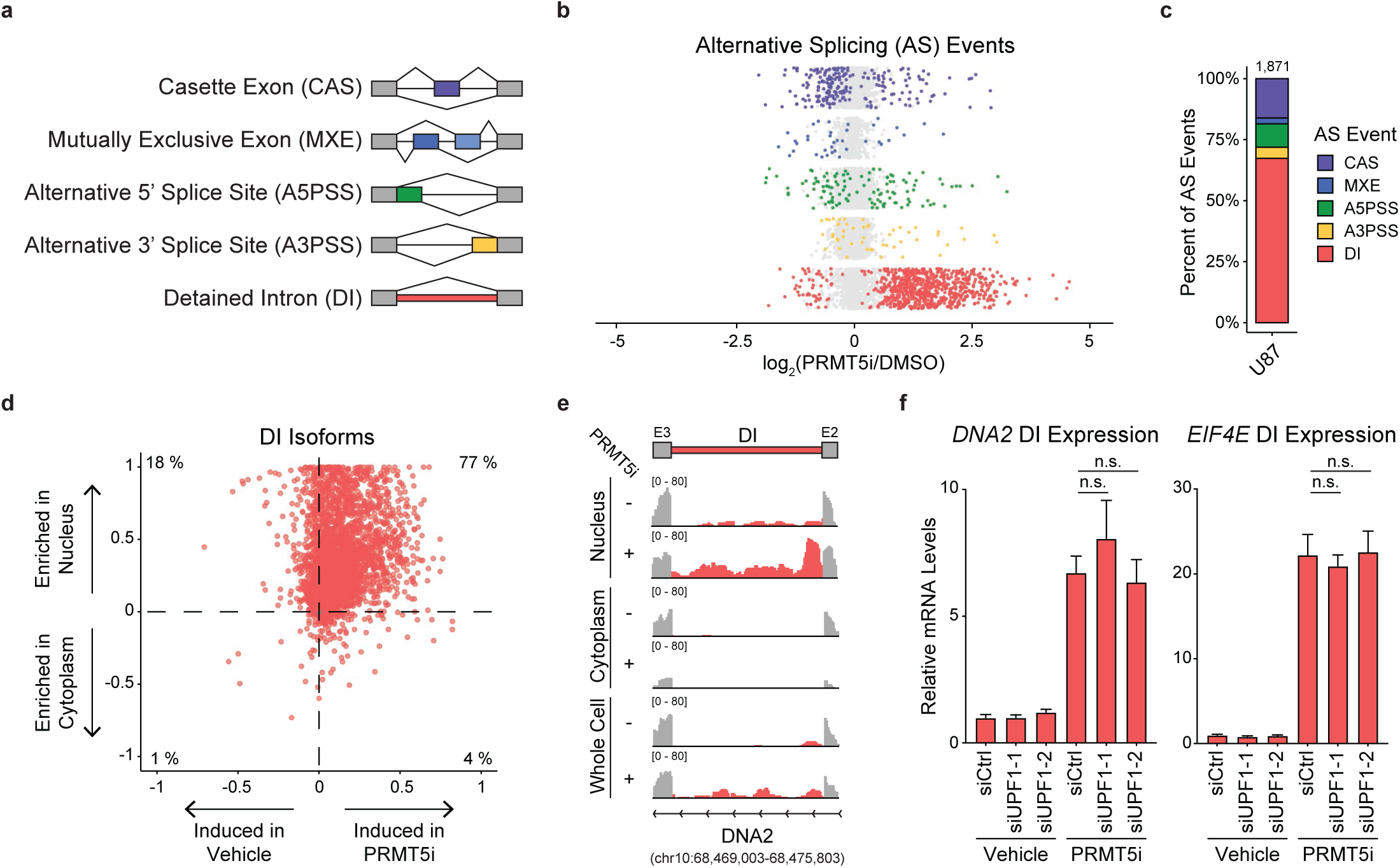
PRMT5 inhibition causes formation of DIs specifically in the nucleus. a-c. U87 cells were treated with vehicle or 10 nM JNJ-64619178 (PRMT5i) for 3 days and polyA+ mRNA was deep sequenced and analyzed to identify and quantify exon-based alternative splicing (AS) events, including DIs, as shown in the schematic (**a**). **b.** Dots represent individual events for each AS type, with color denoting ones that are significantly (p_adj_<0.05, color) or not significantly (gray) altered by treatment. **c.** Proportion of each AS event type that is significantly (p_adj_<0.05) altered by treatment. Number above bar shows total number of significant events. **d.** Quantification of DI isoform induction levels and sub-cellular localization after 3-day vehicle or 10 nM PRMT5i treatment followed by cellular fractionation. Each dot represents a specific DI-containing isoform, where the x-axis value represents the proportion of reads in the vehicle versus PRMT5i-treated samples and the y-axis value represents the proportion of reads in the nuclear versus cytoplasmic fraction. **e.** Expression tracks of the DI in *DNA2*, along with its adjacent exons, present in whole cell, nuclear or cytoplasmic fractions from U87 cells after 3-day vehicle or 10 nM PRMT5i treatment. Red regions indicate intronic reads and gray regions exonic reads. **f.** Relative abundance of *DNA2* (*Left*) or *EIF4E* (*Right*) DIs in U87 cells after 3-day vehicle or 10 nM PRMT5i treatment that were also transfected with two individual siRNAs targeting UPF1 or a non-targeting siRNA for 2 days prior to harvest. p=n.s., Student’s t-test.

We conducted deep sequencing and quantified the levels of coding (fully-spliced) versus intron-containing isoforms present in the cytoplasmic and nuclear compartments (**Figure S1a**). The vast majority of coding isoforms were detected in both compartments (**Figure S1c-d**). In contrast, intron-containing isoforms showed profound nuclear enrichment (**Figure S1c-d**). Indeed, in most cases their cytoplasmic counts were essentially zero. This is consistent with these being DIs, and not RIs. We note that DIs are present in the absence of PRMT5 inhibition, befitting their known occurrence at low levels in normal cells^21^, but their levels dramatically increase with PRMT5i treatment (**Figure 1d-e**).

A second feature that distinguishes DIs from RIs is that DIs are not subject to NMD. Thus, as an additional test, we treated U87 cells with 10 nM PRMT5i for 3 days, and introduced siRNAs against UPF1, a critical NMD-initiating factor, or a non-targeting control, for the last 2 days of treatment. We confirmed siRNA-mediated UPF1 knockdown through RT-qPCR and western blotting (**Figure S1e-f**, p<0.01), and then quantified the levels of DIs in two genes, *DNA2* and *EIF4E* (**Figure 1f**). In both cases, DI levels increase with PRMT5i treatment (p<0.05) and were unaffected by depletion of the NMD machinery (p=n.s.), reinforcing that PRMT5 inhibition specifically regulates the formation of DIs.

### DIs are highly conserved across human cell contexts

Having demonstrated that DI formation is the major splicing change in response to PRMT5 inhibition in U87 cells, we wanted to determine if this is conserved across different human cell lines. For this analysis, we examined six additional lines, including four different cancers, a virally-transformed cell line, and an immortalized normal cell line (**Figure 2a**). We treated these with 10 nM PRMT5i for 3 days and generated whole cell extracts to avoid differences in recovery that might result from unique cellular fractionation conditions (**Figure S2a-b**). Deep sequencing and AS analysis showed that DIs accounted for the majority (62-75%) of AS events significantly altered in response to PRMT5 inhibition in every context (**Figure 2b** and **Figure S2c**; p_adj_<0.05). We identified 2,794 unique DIs that were significantly altered in response to PRMT5 inhibition in at least one of the cell lines (**Figure 2c**; p_adj_<0.05). The vast majority (87-98%) of these were upregulated in response to drug treatment (**Figure 2c** and **Figure S2c**).

**Figure 2.**
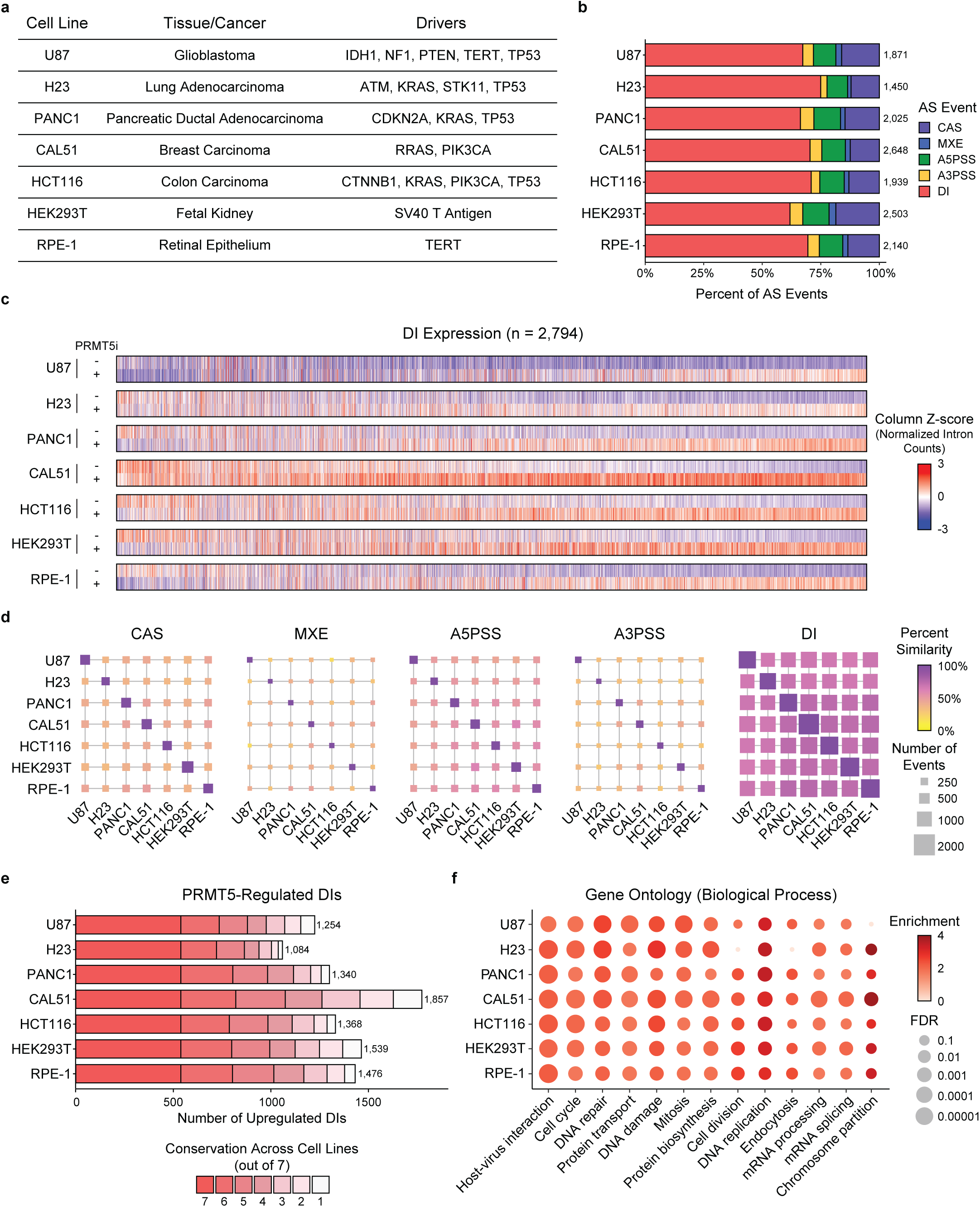
PRMT5 regulates a conserved set of DIs across multiple human cell lines. **a.** Human cell lines used for AS analysis, including cancer/tissue of origin and driver mutations/constructs. **b.** Proportion of each AS event type that show significantly altered levels in the indicated cell line in response to 3-day treatment with 10 nM JNJ-64619178 (PRMT5i) compared to vehicle control (p_adj_<0.05). Number to the right of each bar represents the total number of significant AS events for that cell line. **c.** Relative expression of individual DIs across all human cell lines after 3-day vehicle or PRMT5i treatment. Each column represents a specific DI that is significantly up-or down-regulated by PRMT5i-treatment in at least one cell line. Coloring represents Z-score normalized intron counts, where red indicates higher relative expression and blue indicates lower relative expression. **d.** Similarity matrix of the indicated class of AS event that was significantly altered by PRMT5i treatment. Each square represents the similarity between the two intersecting human cell lines, with coloring indicating percent similarity (Dice Similarity Score) and square size indicating the number of similar events. **e.** Bar charts of DI conservation across the 7 tested human cell lines. The bar length represents the number of significant PRMT5i-upregulated DIs (p_adj_<0.05, log_2_FC>0) for each cell line and the color denotes the number of cell lines in which a given DI is conserved across. The number to the right of the bar represents the total number of significant DIs in each line. **f.** Enriched GO terms from the KW Biological Process gene set for significantly PRMT5i-upregulated DIs in each cell line (p_adj_<0.05, log_2_FC>0). Terms are displayed if they are significant in at least one cell line (FDR<0.01). Color represents enrichment score, while size inversely correlates with significance value.

Having demonstrated that PRMT5 inhibition drives robust DI formation, we then investigated the degree to which DI identity is conserved in different cells. Initially, we performed pairwise comparisons of PRMT5i-regulated AS events, including exon-based AS events and DIs, between all seven cell lines (**Figure 2d**; p_adj_<0.05). For each AS class, we plotted the degree of pairwise conservation as well as the absolute number of conserved events (**Figure 2d**). This revealed remarkably little conservation for any of the exon-based AS events (**Figure 2d**). In contrast, the DI identity was highly conserved for all pairwise comparisons, affecting more than 1,000 individual DIs (**Figure 2d**). Given this high degree of pairwise conservation, we then considered the overlap across all of the lines (**Figure 2e** and **Figure S2d**). This identified a strong degree of conservation amongst PRMT5i-regulated DIs; more than 500 were significantly upregulated in 7/7 cell lines, and more than 50% in any one cell line were conserved in at least 5/7 lines (**Figure 2e** and **Figure S2d**; p_adj_<0.05, log_2_FC>0). The level of conservation was similar in the transformed lines and normal RPE-1 cell line (**Figure 2b,e**). These data indicate that PRMT5 inhibition upregulates a common set of DIs, irrespective of tissue type, driving mutation, or oncogenic status.

Given the high degree of conservation, we next determined the biological processes that PRMT5-regulated DIs are associated with. Specifically, we performed gene ontology (GO) analysis for all upregulated DIs in each line (p_adj_<0.05, log_2_FC>0). In each cell line, there was strong enrichment of core proliferation-associated processes, including multiple cell cycle-associated gene sets (**Figure 2f**), similar to our prior analysis of U87 cells^2^. Together, these data support a model where PRMT5 enables efficient intron splicing of a conserved set of DIs in proliferation-associated genes.

### DI upregulation from methylosome impairment mirrors viability impairment

Our observations are highly consistent with the fact that PRMT5i causes proliferation defects, as well as the prevailing view that splicing defects underlie the vulnerability of cancer cells to PRMT5i. However, neither our, nor existing, data directly address the importance of the PRMT5-splicing axis because they utilize inhibitors that disrupt all PRMT5 functions (**Figure 3a**). To address this limitation, we wanted to specifically interrogate the necessity of the methylosome, which mediates PRMT5’s core splicing role, without disrupting other PRMT5 functions. To achieve this, we used the dTag degron system^26^ to specifically deplete CLNS1A and thus methylosome function (**Figure 3a**). We selected three cell lines – CAL51, HEK293T, and HCT116 (**Figure 2a**) – and introduced the dTag degron domain, which also contains an HA-tag, into the endogenous *CLNS1A* locus. For each line, we identified four clones that have homozygous degron cassette insertion, as determined by a 12 kDa molecular weight increase for CLNS1A and acquired HA positivity (**Figure 3b** and **Figure S3a**). The degron inducer molecule, dTag-13, has off-target toxicity at high concentrations; thus, we challenged parental cell lines with a range of dTag-13 doses for 6 days and showed that doses up to 1 μΜ avoided toxicity (**Figure 3c**). We then treated all of our clones for 3 days with 1 μΜ dTag-13 and showed that this achieved high levels of CLNS1A depletion for all cell lines (**Figure 3d**).

**Figure 3.**
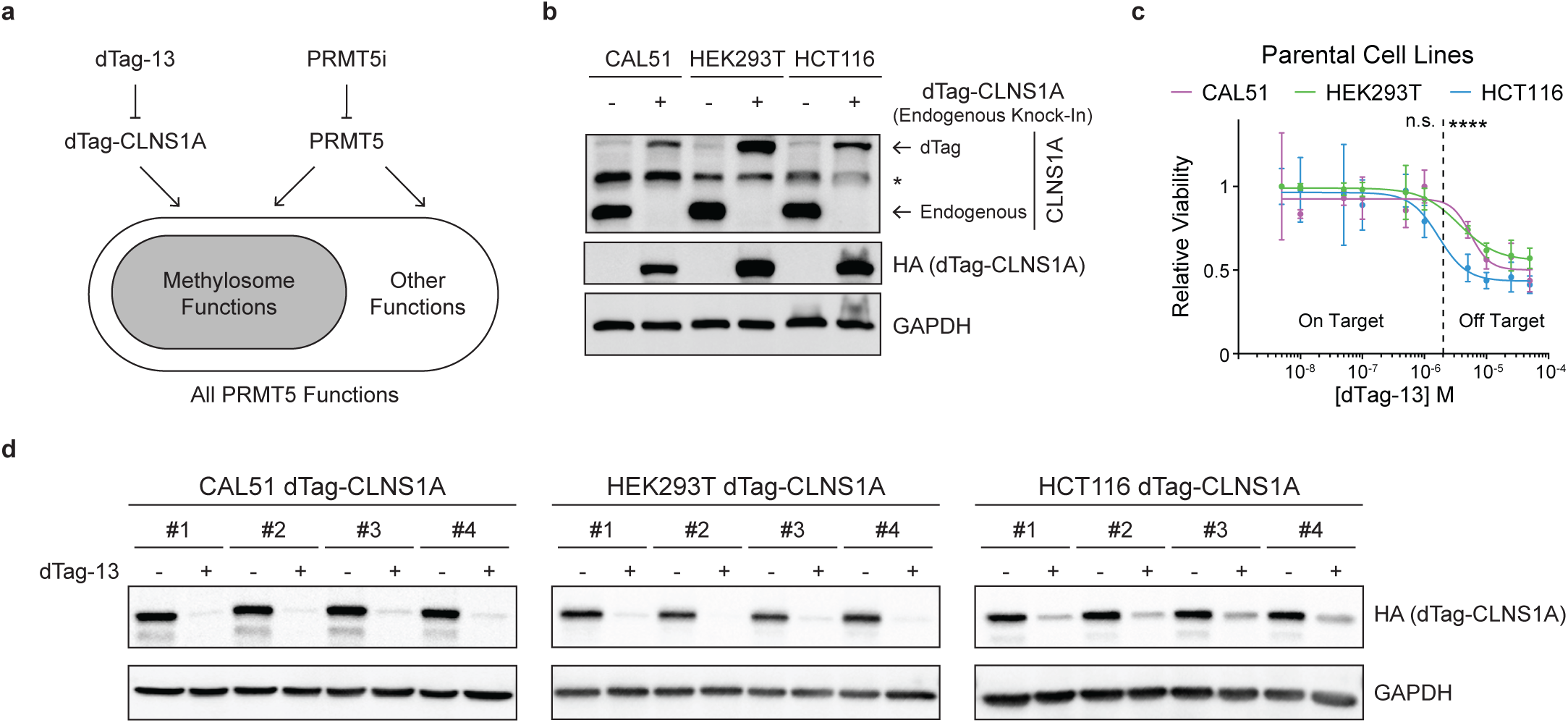
Generation of endogenous dTag-CLNS1A in multiple human cell lines to specifically inhibit the PRMT5-splicing axis. **a.** Schematic indicating how use of the dTag-CLNS1A degron system specifically separates disruption of the methylosome from other PRMT5 functions. **b.** Western blot analyses of representative dTag-CLNS1A clones for CAL51, HEK293T, and HCT116 cells, alongside parental controls with antibodies against CLNS1A, the inserted HA tag, or loading control GAPDH. Asterisk (*) marks a non-specific band. **c.** Dose response curves for three parental cell lines treated with dTag-13 for 6 days. Data are mean ± SD of 3 technical replicates/line. Indicated p-values represent the most significant comparison between the respective drug concentration and vehicle-treated cells. Vertical dashed line represents the concentration above which dTag-13 has off-target effects. ****p<0.0001, Student’s t-test. **d.** Western blot analyses of HA-CLNS1A and loading control GAPDH in four CAL51, HEK293T, and HCT116 dTag-CLNS1A clones treated with either vehicle or 1 μM dTag-13 for 3 days.

Having validated our dTag-CLNS1A clones we then compared their sensitivity to CLNS1A depletion versus PRMT5 inhibition. Specifically, we treated our clones with vehicle, 1 μM dTag-13, or 10 nΜ PRMT5i, harvesting after 3 days for molecular analyses or 6 days for viability. PRMT5i behaved as expected; at a molecular level, it had no detectable effect on the levels of HA-CLNS1A (**Figure 4a-b**; p=n.s.) but caused a dramatic downregulation in the methylation of SmB proteins (**Figure 4a-b**; p<0.01), as well as all other SDMA-modified proteins (**Figure S4a**). These molecular changes corresponded to a loss of cellular viability, with similar IC50 values for all clones (**Figure S3b**). We then examined the consequences of dTag-13 treatment. Consistent with our clone validation (**Figure 3d**), we saw robust depletion of HA-CLNS1A in all clones (**Figure 4a-b**; p<0.01). Unexpectedly, we observed two distinct downstream phenotypes, which depended on the cell line. CAL51 and HEK293T displayed one set of phenotypes. First, dTag-13 treated clones showed specific loss of methylation at SmB (**Figure 4a-b**; p<0.05) but not other PRMT5 targets (**Figure S4a**). Second, they displayed robust DI induction (**Figure 4c**), alongside far fewer significant changes in exon-based AS events (**Figure 4c** and **Figure S4c**). There was a high degree of overlap between DIs affected by CLNS1A-deficiency versus PRMT5i treatment in the relevant parental CAL51 or HEK293T lines, while other AS events showed less overlap between these treatments (**Figure 4d** and **Figure S4d-g)**. Finally, dTag-13 treated clones showed a profound loss of cellular viability for CAL51 and HEK293T (**Figure 4e**; p<0.001). The HCT116 clones displayed starkly different phenotypes in response to dTag-13; there was no significant loss of Sm protein methylation (**Figure 4a-b**; p=n.s.), very little significant induction of DIs or other AS changes (**Figure 4c-d** and **Figure S4c-g**), and no significant loss of cellular viability (**Figure 4e**; p=n.s.). Collectively, these data argue that PRMT5-mediated Sm protein methylation is critical for maintenance of normal splicing fidelity – particularly limiting DI levels – in some cell lines, and disruption of this axis represents a critical vulnerability in sensitive cancer cells.

**Figure 4.**
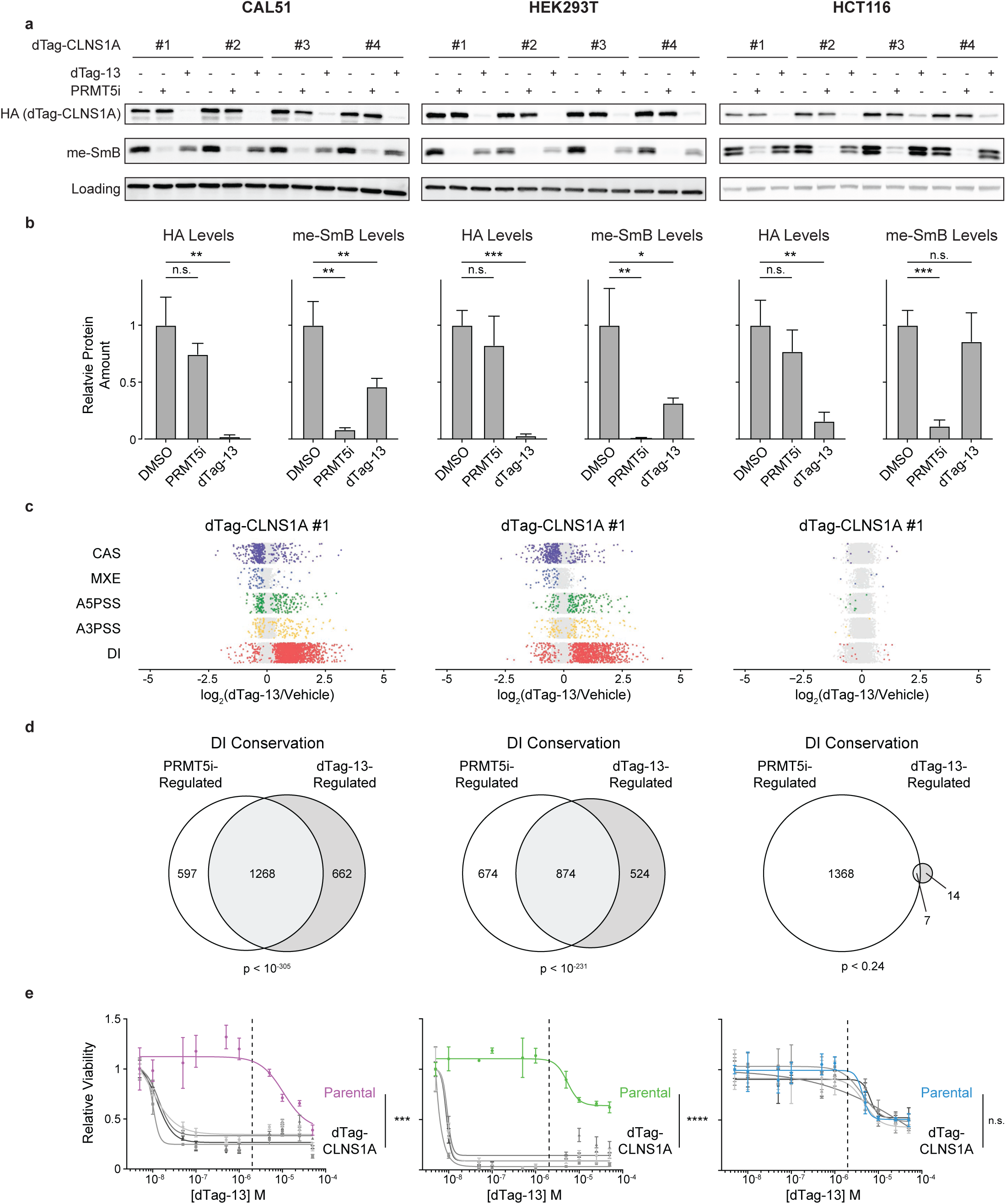
Loss of Sm protein methylation correlates with increased DI formation and decreased viability. **a.** Western blot analyses of HA-CLNS1A, methyl-SmB, and GAPDH (CAL51 and HEK293T) or HSP90 (HCT116) as a loading control in four CAL51, HEK293T, and HCT116 dTag-CLNS1A clones treated with either vehicle, 10 nM JNJ-64619178 (PRMT5i), or 1 μM dTag-13 for 3 days. **b.** Quantification of HA (*Left*) and methyl-SmB (*Right*) levels from **a,** normalized to loading control. Data are mean ± SD of 4 biological replicates. *p<0.05, **p<0.01, and ***p<0.001, Welch’s t-test. **c.** Quantification of indicated classes of AS event in dTag-CLNS1A #1 for CAL51, HEK293T, and HCT116 treated with vehicle or 1 μΜ dTag-13 for 3 days. Dots represent individual events, denoting ones that are significantly (p_adj_<0.05, color), or not significantly (gray), altered by dTag-13 treatment. **d.** Overlap of DIs in CAL51, HEK293T, and HCT116 cells that were significantly altered by dTag-13-induced CLNS1A depletion versus PRMT5i treatment, compared to their vehicle controls (p_adj_<0.05). Overlap significance was calculated using a hypergeometric test. **e.** Dose response curves of four dTag-CLNS1A clones versus parental controls for the indicated CAL51, HEK293T, and HCT116 lines after a 6-day treatment with dTag-13. Data represents mean ± SD of 3 technical replicates. ***p<0.001 and ****p<0.0001, Student’s t-test.

### CLNS1A is fully dispensable in some contexts

The finding that all HCT116 dTag-CLNS1A clones were insensitive to dTag-13 treatment was highly unexpected, given that CLNS1A, and the methylosome, are described as cell essential. We realized that dTag-13 does not achieve complete CLNS1A loss and the residual levels in HCT116 are typically higher than those observed in CAL51 or HEK293T clones (**Figure 4a-b**). Thus, we wondered whether a more prolonged dTag-13 treatment would reveal an impairment in HCT116. To address this, we treated two HCT116 dTag-CLNS1A clones with 1 μM dTag-13 for 50 days, harvesting cells every 5 days for replating. These cells showed no perceptible growth impairment and no noticeable loss of SmB methylation, even though CLNS1A was efficiently depleted throughout the time-course (**Figure 5a**). Thus, HCT116 can tolerate a dramatic, long-term reduction CLNS1A depletion.

**Figure 5.**
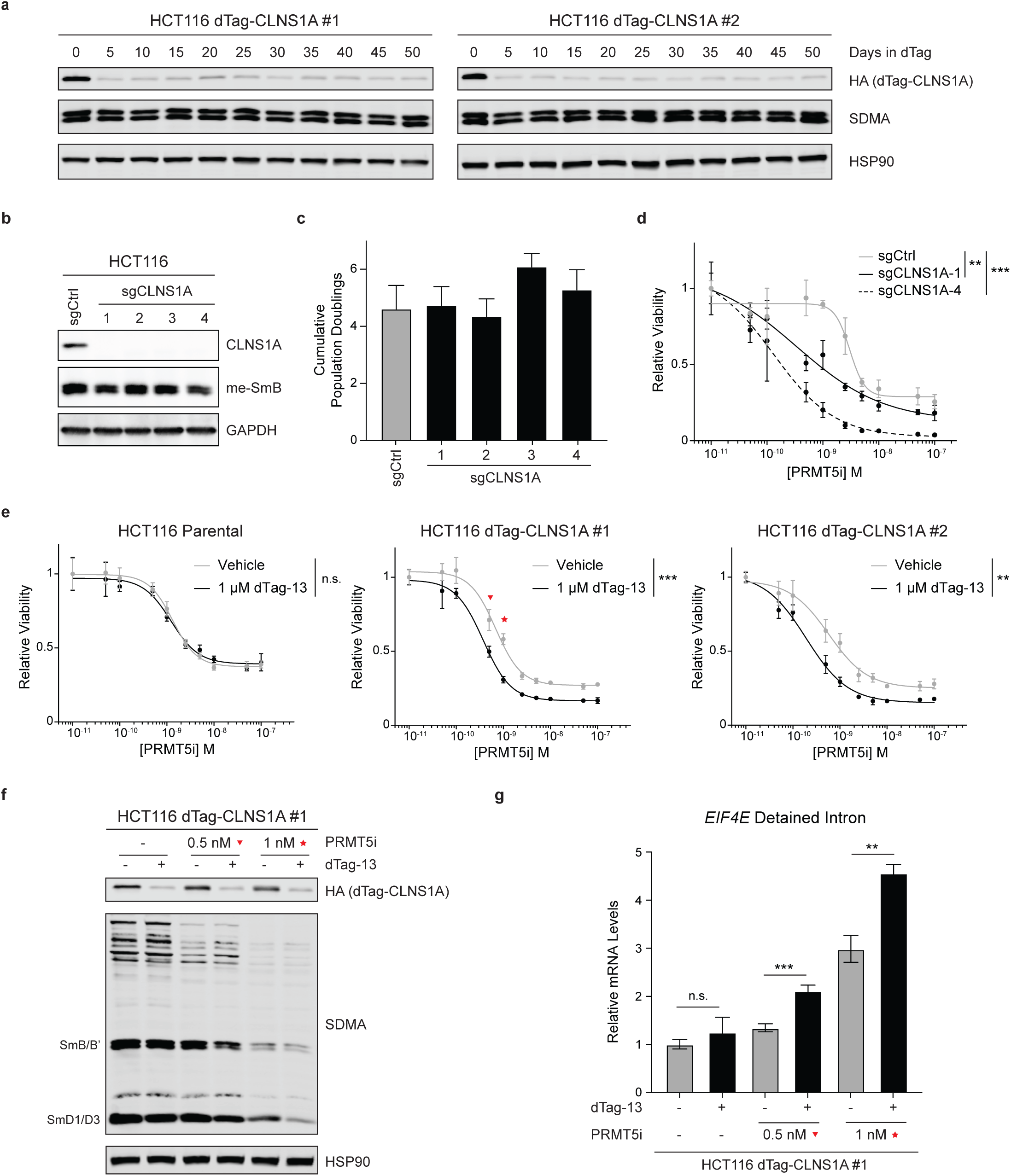
HCT116 cells are CLNS1A-independent until PRMT5 activity is decreased. **a.** Western blot analyses of HA-CLNSIA, methyl-SmB, and HSP90 as loading control in HCT116 dTag-CLNS1A #1 (*Left*) and #2 (*Right*) maintained in 1 μM dTag-13 for the indicated number of days. **b.** Representative western blot analysis of CLNS1A, methyl-SmB, and loading control GAPDH levels in HCT116 sgCtrl and four sgCLNS1A clones (n=4 biological replicates). **c.** Relative proliferation of HCT116 sgCtrl and four sgCLNS1A clones through analyses of cumulative population doubling over a 6-day period. Data represents mean ± SD of 10 technical replicates. **d.** Dose response curve of HCT116 sgCtrl and indicated representative sgCLNS1A clones treated with the PRMT5i JNJ-64619178 for 6 days. Data represents mean ± SD of 3 technical replicates. **p<0.01 and ***p<0.001, Student’s t-test. **e.** Dose response curve of HCT116 parental or two dTag-CLNS1A clones treated with or without 1µM dTag-13 in combination with the indicated concentrations of JNJ-64619178 (PRMT5i) for 6 days. Data represents mean ± SD of 3 technical replicates. **p<0.01 and ***p<0.001, Student’s t-test. **f.** Western blot to assess levels of HA-tag, SDMA, and loading control HSP90 in extracts from HCT116 dTag-CLNS1A #1 after combination treatment with the indicated concentrations of PRMT5i and dTag-13 for 3 days. Red markings correspond to the dose of PRMT5i used in **Figure 5e**. **g.** Relative levels of a representative DI in *EIF4E* in HCT116 dTag-CLNS1A #1 after combination treatment with the indicated concentrations of PRMT5i and dTag-13 for 3 days. Red markings correspond to the dose of PRMT5i used in **Figure 5e**. **p<0.01 and ****p<0.0001, Student’s t-test.

Given this finding, we wondered whether CLNS1A is fully dispensable in HCT116 cells. To address this, we utilized CRISPR-Cas9-mediated gene targeting to attempt to generate *CLNS1A* knock-out lines. We identified two independent knock-out lines (sgCLNS1A-1 and-2), carrying different homozygous deletions that each caused frameshift mutations and premature termination codons (PTCs) (**Figure S5a**). We also generated two independent hypomorphic clones (sgCLNS1A-3 and sgCLNS1A-4), in which one allele bore a small in-frame deletion, Δ117-118, and the other was disrupted by different frameshift mutations and consequent PTCs (**Figure S5a**). Western blotting confirmed that the KO clones lacked detectable CLNS1A and revealed that the hypomorphic clones both displayed effectively undetectable protein levels (**Figure 5b** and **Figure S5b**), suggesting that the in-frame deletion impairs protein stability. We then probed for Sm protein methylation. Remarkably, the HCT116 sgCLNS1A clones retained Sm protein methylation at relatively normal levels in two clones and with only a modest reduction in two others (**Figure 5b** and **Figure S5b**). Most importantly, we found that all four sgCLNS1A clones proliferated at the same rate as the sgCtrl cells (**Figure 5c**).

These data argue that HCT116 cells can thrive in the absence of CLNS1A. Given how unexpected this finding was, we wanted to thoroughly consider other possibilities. We realized that there is an annotated alternatively-spliced *CLNS1A* isoform and that the exon targeted by our sgRNA is not part of the predicted truncated transcript^27^. Thus, we performed RT-PCR for both full length (FL)-and truncated-*CLNS1A* transcripts. We found that both KO clones had significantly reduced levels of FL-*CLNS1A* mRNA and low levels of truncated-*CLNS1A*, which was nearly absent from sgCtrl (**Figure S5c-d**). Thus, we considered whether truncated-*CLNS1A* might be playing a role in the HCT116 KO cells. Initially, we used Alphafold2^28,29^ to model the predicted protein structure (**Figure S5e**). The portion encoded by the skipped exons (shown in red) includes acidic residues (shown as sticks) that are critical ionic contact points between CLNS1A and the basic Sm proteins, as well as a central α-helix and β-sheet in the globular domain that likely enhance protein stability. To directly test this, we infected sgCLNS1A-1 cells with retroviruses encoding FL- or truncated-CLNS1A at a similar multiplicity of infection (MOI) and showed that truncated-CLNS1A is expressed at lower levels than the FL species (**Figure S5f**).

Since CLNS1A functions in a complex with PRMT5, we anticipated that CLNS1A deficiency, and subsequent ectopic expression, would modulate the sensitivity of the HCT116 cells to PRMT5i. Initially, we treated two representative sgCLNS1A lines (sgCLNS1A-1 and-4) with PRMT5i for 6 days, alongside the parental HCT116, and showed that the KO clones showed significantly increased susceptibility to PRMT5i (**Figure 5d**; p<0.01). We then examined the impact of ectopic expression of FL-or truncated-CLNS1A on this response. Compared to empty vector, FL-CLNS1A made all four of the sgCLNS1A clones more resistant to PRMT5i (**Figure S5g-h**; p<0.01), while truncated-CLNS1A had no effect in the sgCLNS1A-1 cells (**Figure S5i**; p=n.s.). These data yield two major conclusions. First, they show that truncated-CLNS1A is non-functional, reinforcing the notion that the sgCLNS1A clones are CLNS1A deficient. Second, they show that CLSN1A is fully dispensable for the viability of HCT116 cells, but its loss increases their vulnerability to PRMT5 inhibition.

We hypothesized that we could exploit the synergy between CLNS1A deficiency and PRMT5 inhibition to definitively address the relationship between the PRMT5-splicing axis and cellular viability. For this, we cultured two different HCT116 dTag-CLNS1A clones, alongside the parental line, with either vehicle control or 1 μM dTag-13 across a range of concentrations for two mechanistically distinct PRMT5i [JNJ-64619178 (**Figure 5e**) and EPZ015666^4^ (**Figure S5j**)]. dTag-13 treatment did not affect the responsiveness of the parental HCT116 to either PRMT5i (p=n.s.), as expected, but significantly increased the sensitivity to both PRMT5i in the dTag-CLNS1A lines (p<0.01). We noted that the largest viability differences with or without dTag-13 treatment were observed at 0.5 and 1 nΜ of the PRMT5i JNJ-64619178 (**Figure 5e** *Center*, red triangle and star). Thus, we focused on these concentrations to determine whether differences in Sm protein methylation and DI levels were associated with the observed viability differences. At both 0.5 and 1 nΜ PRMT5i, HCT116 dTag-CLNS1A #1 exhibited a greater loss of Sm protein methylation when dTag-13 treatment was combined with PRMT5i (**Figure 5f**). Moreover, RT-qPCR showed that decreasing Sm protein methylation was accompanied by a significant increase in the levels of the canonical DI in *EIF4E* (**Figure 5g**; p<0.01). Thus, while HCT116 dTag-CLNS1A clones cells can survive in the absence of CLNS1A, this depletion significantly sensitizes cells to PRMT5 inhibition, which correlates with increased impairment of Sm protein methylation and significantly increased DI formation.

### MTAP deficiency increases dependence on methylosome activity

We also considered how the status of *methylthioadenosine phosphorylase* (*MTAP*) modulates the influence of the methylosome and DI regulation on cell viability. *MTAP* is frequently co-deleted in many human tumors due to its adjacency to the *CDKN2A* tumor suppressor locus^30–35^. This causes accumulation of the MTAP substrate, MTA, which serves as an endogenous inhibitor of PRMT5, potentially increasing sensitivity to PRMT5i^36,37^. If DIs represent the core vulnerability of cancer cells, then MTAP loss should synergize with CLNSIA deficiency in our HCT116 dTag-CLNS1A clones. To address this, we used CRISPR-Cas9-mediated gene targeting to generate two independent *MTAP*-deficient variants of HCT116 dTag-CLNS1A #1 (**Figure S5k**). We then treated sgCtrl and sgMTAP lines with vehicle or 1 μM dTag-13 for 6 days and assessed cellular viability. The MTAP-deficient HCT116 CLNS1A-dTag #1 cells showed reduced viability when treated with 1 μM dTag-13, in contrast to the MTAP-proficient sgCtrl (**Figure S5k**). These results confirm that *MTAP*-deficiency sensitizes cells to inhibition of the PRMT5-splicing axis, and highlights the importance of studying PRMT5 biology in both MTAP-proficient and deficient contexts. They also reinforce our conclusion that maintenance of appropriate splicing, especially DI levels, is critical in enabling cellular viability.

### DI identity is conserved across mouse cell lines to regulate proliferation-associated processes

Our data show that DIs are highly conserved across human cell types and support that they are the key vulnerability to PRMT5 inhibition. Given this, we expanded our analysis to murine cells. Specifically, we treated murine lung and pancreatic ductal adenocarcinoma lines for 3 days with PRMT5i [JNJ-64619178 (100 nM) or EPZ015666 (10 μM)] (**Figure 6a** and **Figure S6a**). We performed deep RNA-sequencing and conducted AS analysis, which showed that DIs accounted for the majority of significant AS events irrespective of cell line or inhibitor (**Figure 6b** and **Figure S6c**; p_adj_<0.05). Moreover, DI identity was highly conserved across murine cell lines (**Figure 6c-e**), and much more so than exon-based AS events (**Figure 6d**). Indeed, nearly 1000 individual DIs were common to all murine cell lines examined (**Figure 6e**). Thus, similar to our finding in human cells (**Figure 2c-e**), these data show that PRMT5 regulates a conserved set of murine DIs. We performed GO analysis on all upregulated DIs for each murine line (p_adj_<0.05, log_2_FC>0) and observed strong enrichment for core proliferation-associated processes (**Figure 6f**), including multiple cell cycle-associated gene sets, common to all lines. Collectively, our data show that PRMT5 inhibition drives the formation of a conserved set of DIs in murine cells that impacts proliferation-associated processes.

**Figure 6.**
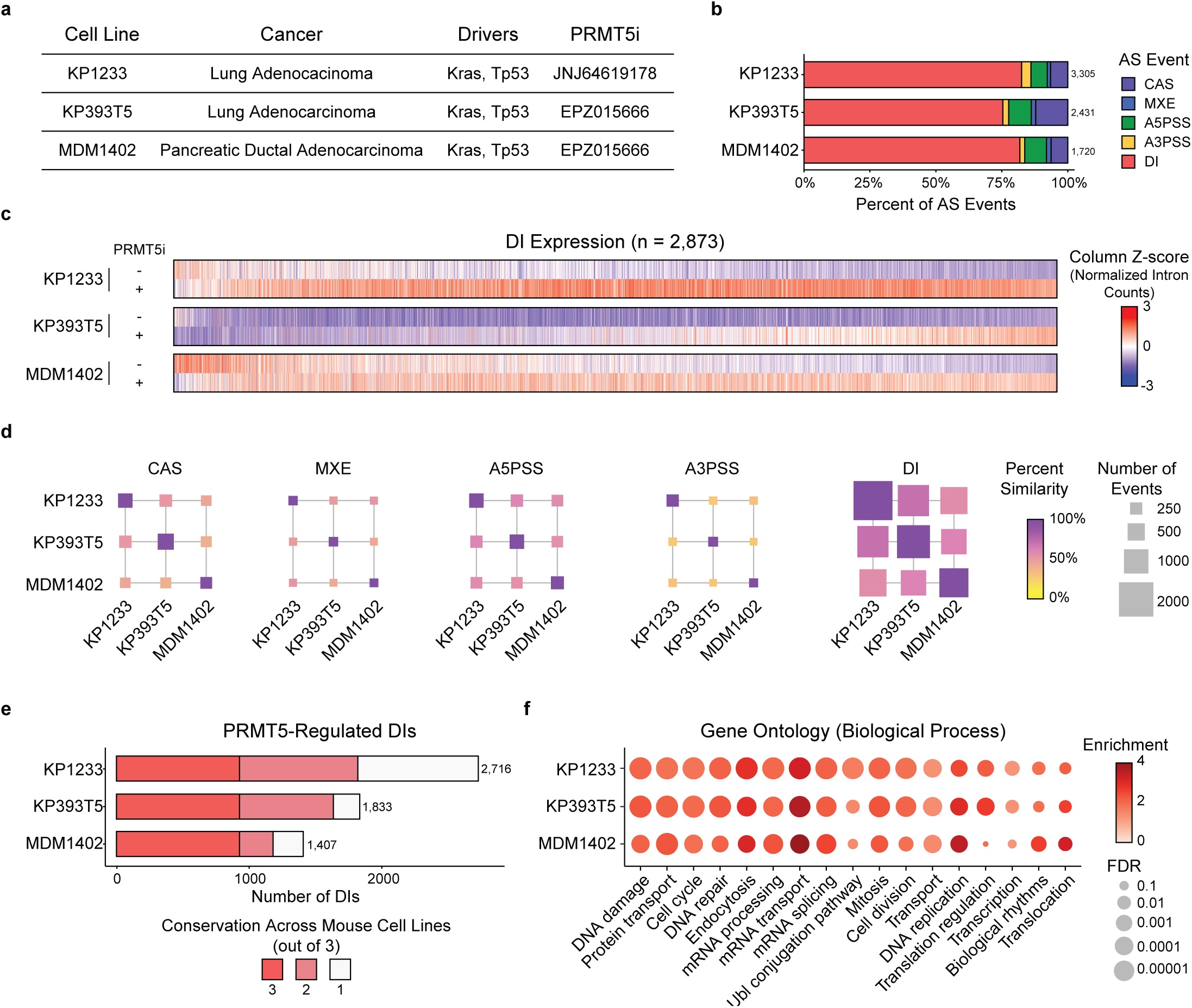
PRMT5 regulates a conserved set of DIs in multiple mouse cell lines. **a.** Summary of mouse cell lines used for AS analysis, including cancer type, driver mutations, and PRMT5i used. **b.** Proportion of each AS event type that is significantly different in indicated cell line after 3-day PRMT5i treatment compared to vehicle control (p_adj_<0.05). Number above bar represents the total number of significant AS events. **c.** Relative expression of individual DIs across all mouse cell lines after 3-day vehicle or PRMT5i treatment. Each column represents a specific DI that is significantly upregulated or downregulated by PRMT5i-treatment in at least one cell line. Colors represent column Z-score normalized intron counts, where red indicates higher relative expression and blue indicates lower relative expression. **d.** Similarity matrix of the indicated class of AS event that was significantly altered by PRMT5i treatment. Each square represents the similarity between the two intersecting murine cell lines. Square coloring indicates percent similarity (Dice Similarity Score), and square size is proportional to the number of similar events. **e.** Bar charts of DI conservation across all mouse cell lines. Each bar represents the number of PRMT5i-upregulated DIs (p_adj_<0.05, log_2_FC>0) in each cell line and the color corresponds to the number of cell lines in which a given DI is conserved across. Number to the right of the bar represents the total number of significant DIs in each line. **f.** Enriched GO terms from the KW Biological Process gene set for PRMT5i-upregualted DIs in each cell line (p_adj_<0.05, log_2_FC>0). Terms are displayed if they are significant in at least one cell line (FDR<0.01). Color represents enrichment score, while size inversely correlates with significance value.

### DIs are not conserved across species, but the biological complexes and processes they affect are highly conserved

Our analyses indicate that DIs impinge on similar biological processes in both human and mouse cells (**Figure 2f** and **Figure 6f**). To explore this overlap more closely, we conducted pairwise comparisons of the processes affected by PRMT5i-induced DIs in the human H23 lung cancer line with those detected in each of the murine lung cancer lines, KP393T5 and KP1233, and observed 71% (**Figure 7a-b**) and 79% (**Figure S7a**) overlap, respectively. Again, there was high representation of core proliferation-associated processes. Similarly high conservation (56%) was observed when comparing human PANC1 and murine MDM1402 pancreatic cancer lines (**Figure S7b**). This high inter-species conservation of DI-affected biological processes was also agnostic of tissue type, as comparison of the biological processes conserved across all human lines with those conserved across all murine lines showed 67% conservation (**Figure S7c**).

**Figure 7.**
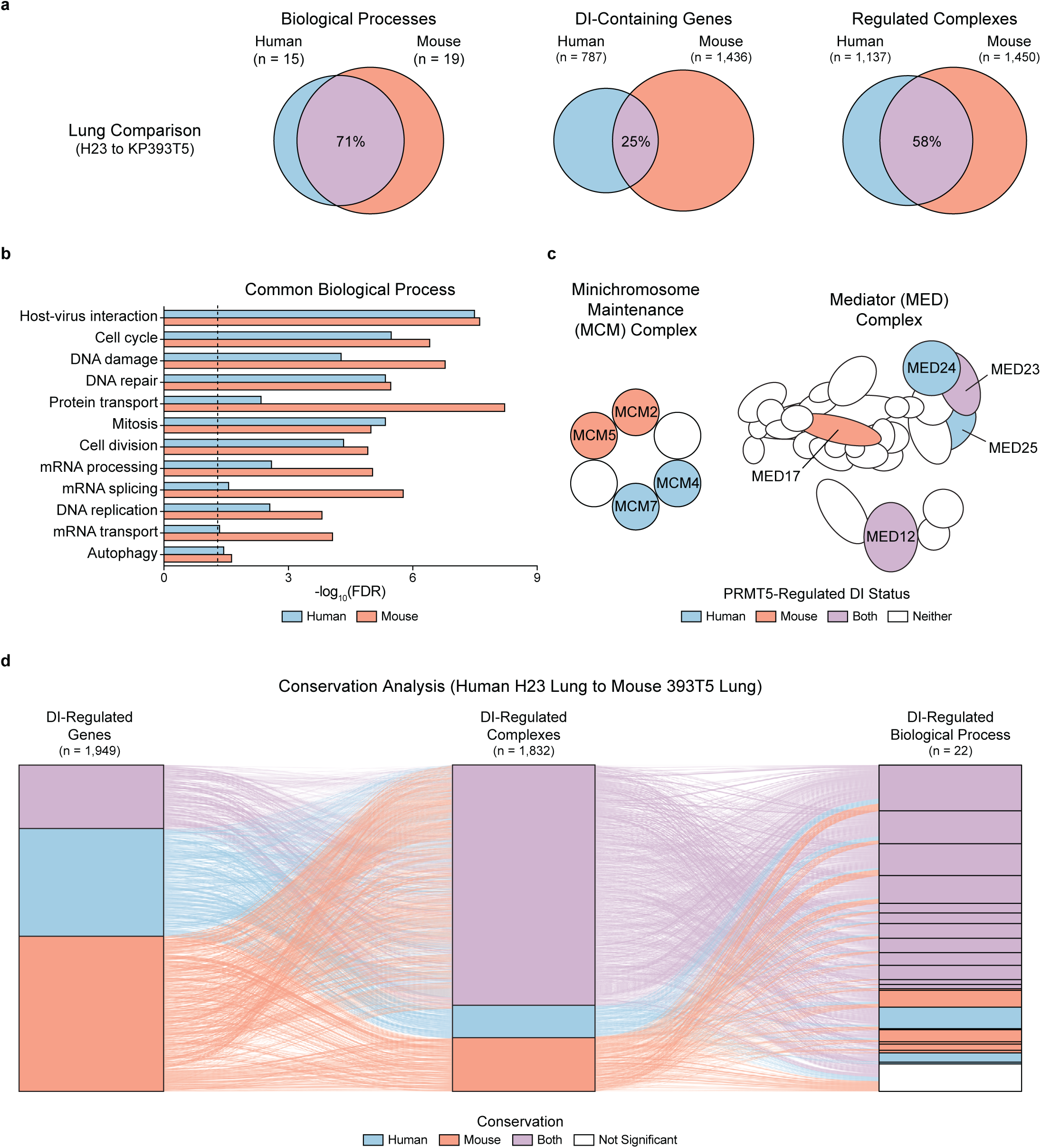
PRMT5-regulated DI identity is not conserved across species, but they occur in genes in similar complexes to regulate proliferation-associated processes. **a.** Overlap of common DI-regulated (*Left*) processes, (*Center*) genes, and (*Right*) complexes for PRMT5-regulated DIs in human (H23) and mouse (KP393T5) lung cancer cell lines. Percentages represent the Dice Similarity Score for each comparison. **b.** Enrichment of DI-regulated biological processes in human and mouse cells. Each pair of bars represents a specific biological process, with the blue bar representing the FDR of the human H23 lung cancer line and the red bar representing the FDR of the mouse KP393T5 lung cancer line. Vertical dashed line represents the significance cutoff (FDR < 0.05). **c.** Schematic of representative DI-regulated complexes, including the minichromosome maintenance (MCM) and mediator (MED) complexes. Each shape represents a specific complex member, and a color fill of blue, red, purple, or white indicates that the complex member contained a DI in human, mouse, both species, or neither, respectively. **d.** Sankey plot analysis of convergence of DI-regulated complexes and biological processes. Each line represents a specific DI moving from (*Column 1*) gene to (*Column 2*) complexes to (*Column 3*) biological processes. Line color signifies conservation at the (*left*) gene or (*right*) complex level, and thickness corresponds to number of (*Left*) complexes or (*Right*) biological processes a given gene is a part of. Column color represents conservation at the level of (*Column 1*) gene, (*Column 2*) complexes, and (*Column 3*) biological processes. Blue indicates unique to human cells, red unique to mouse cells, and purple common to both species. For biological processes, white indicates DIs/complexes that map to non-significant biological processes (FDR>0.05).

We anticipated that the inter-species overlap would reflect high conservation of the DI-containing genes. However, we observed a striking lack of inter-species conservation in DI-containing genes, even when we considered a specific tissue type (21%-25%; **Figure 7a** and **Figure S7a-b**) or the genes that were commonly-regulated in all human versus all mouse lines (25%; **Figure S7c**). Moreover, when genes were conserved between human and mouse, the DIs typically affected distinct introns. Thus, our data show that PRMT5i-induced DIs are poorly conserved across species at the gene level but somehow impact the same biological processes. As a possible explanation, we realized that many DI-containing genes encode components of multi-protein complexes. Consequently, we performed CORUM analysis^38^ to identify complexes affected by DI-containing genes in mouse versus human cells and then quantified the degree of overlap. For both comparisons within a specific tissue types (**Figure 7a** and **Figure S7a-b**) and comparison of DIs commonly-regulated in all human versus all mouse lines (**Figure S7c**), we found a strong overlap in complex identity (53-59%), which was much higher than the DI-containing gene identity and approached the levels of biological process conservation. Examples of affected complexes are shown in **Figure 7c** and **Figure S7d-e**. Moreover, examination of representative complexes revealed many cases where the PRMT5-regulated DIs impacted more than one complex component within a single species (**Figure 7c** and **Figure S7d-e**), which would further exacerbate defects in the assembly and levels of functional complexes. Finally, analyses at the global level (**Figure 7d** and **Figure S7f**) clearly illustrate how DI upregulation in species-specific genes converges on multi-protein complexes to impact common, fundamental biological processes.

## Discussion

A central goal of this study was to determine which of PRMT5’s many functions account for its oncogenic role and, thus, the cellular vulnerability to PRMT5 inhibitors. The prevailing view in the field is that this reflects PRMT5’s role in splicing. This hypothesis is based on numerous reports that PRMT5 inhibition induces splicing defects; however, this has not been directly tested because these studies used chemical or genetic approaches that disrupt all PRMT5 functions. Our current study selectively perturbs the PRMT5-splicing axis through induced degradation of the Sm protein specific PRMT5 cofactor, CLNS1A. We found that two cell lines were unable to survive CLNS1A depletion, while a third cell line (HCT116) remained fully viable in response to either CLNS1A depletion or complete *CLNS1A* knockout. This latter finding was completely unexpected as *CLNS1A* is widely believed to be pan-cell essential. However, this view is based primarily on large-scale depletion screens, which effectively test fitness rather than essentiality. Based on reports that, at least *in vitro*, PRMT5-MEP50 complexes are sufficient to bind and methylate Sm proteins^22,39^, we speculate that there is enough PRMT5-MEP50 activity in HCT116 cells to survive without CLNS1A. Regardless of the mechanism by which HCT116 cells achieve CLNS1A independence, the differential viability of our three cell lines has helped elucidate the causal events of PRMT5i-mediated tumor suppression. Specifically, both CLNS1A- dependent cell lines exhibited loss of Sm protein methylation and robust induction of splicing defects upon CLNS1A depletion, while the CLNS1A-independent HCT116 cells showed no changes for either phenotype. Importantly, we find that the CLNS1A-deficient HCT116 cells show increased sensitivity to PRMT5i or genetic ablation of *MTAP*, which lowers endogenous PRMT5 activity. The PRMT5i doses that define the differential viability window of HCT116 with or without CLNS1A-depletion correspond exactly to differences in Sm protein methylation and DI levels between these two contexts. Collectively, these data establish that the PRMT5-splicing axis is critical to maintain cellular viability.

What is the critical aspect of splicing? Our data show that upregulation of unspliced introns, which we experimentally validated as *bona fide* DIs, is a defining hallmark of effective PRMT5 inhibition. They further argue that DIs, rather that canonical exon-based AS events, represent the core vulnerability to PRMT5 inhibition. Specifically, DIs are the most prevalent change, they are almost always upregulated, and they show a high level of identity conservation across numerous cell lines within a species. Moreover, the PRMT5i-responsive DIs selectively impact genes that control cell cycle regulation and proliferation, the biological processes that are robustly, and consistently, impacted by effective PRMT5 inhibition. In contrast to DIs, other AS events are a rarer consequence of PRMT5 inhibition and show minimal conservation, even within a species. We note that we are not ruling out a role for specific AS events; indeed, there is compelling evidence that PRMT5 inhibition can drive alternative splicing of *MDM4*, which results in p53 activation^20^. However, we believe that DI formation is the primary driver of PRMT5i sensitivity, through its impact on proliferation-associated genes, with other AS events potentially having contributing roles in a context-dependent manner.

Our study also reveals fascinating similarities, as well as differences, in DI biology between humans and mice. With respect to similarities, we find on the order of 1000-2000 genes that are affected by PRMT5i-responsive DIs in either human or mouse cells. These DIs are remarkably well-conserved in an intra-species manner, for example approximately half of the PRMT5i- responsive DIs were conserved in all 7 human cell lines analyzed. This conservation is agnostic to tissue type, whether the cells are transformed or normal, or the nature of the driving mutations within the cancer cells. As noted above, many of affected genes are cell essential and significantly enriched for roles in cellular proliferation. The presence of DIs causes transcripts to be sequestered in the nucleus, as demonstrated by our cellular fractionation, thereby reducing the levels of appropriately-spliced and exported transcript available for translation^2,21^. It is easy to envisage that the combined effect of reduced transcript levels of hundreds, if not thousands, of core proliferation regulators causes profound challenges for cell cycle progression.

Our data further show that the same core biological processes are affected in mouse and human cells. However, remarkably there is minimal conservation between these two species with regard to the identity of the DIs, or even the affected genes. Instead, our analyses argue that these targets frequently represent distinct, rather than overlapping, components of multi-protein complexes involved in proliferative control. Interestingly, in many cases, we find that more than one component of a given complex are encoded by DI-containing genes. It is fascinating to consider how two species have developed a way to regulate the same biological processes through the presence of DIs in distinct genes.

A major impetus for this study was the fact that numerous PRMT5i are currently in, or approaching, clinical trials without a clear understanding of the root cause of cancer vulnerability. Our study strongly validates the prevailing view that the effectiveness of PRMT5i reflects induction of splicing defects within proliferating cells. To improve cancer cell selectivity, the field has already generated a class of PRMT5i that specifically targets cells that are deficient for *MTAP*, which typically reflects loss of the *CDKN2A* tumor suppressor locus. We hypothesize that PRMT5i that more specifically target the PRMT5-splicing axis, might offer further selectivity and/or efficacy.

## Resource Availability

All data needed to evaluate the conclusions in the paper are present in the paper and/or the Supplementary Materials. RNA-seq data that support the findings of this study will be uploaded to the Gene Expression Omnibus (GEO) prior to publication, and can be provided upon request. All custom code used in this work is available from the corresponding author.

## Acknowledgements

We are especially grateful to our colleague, Phillip A. Sharp, for insightful discussions throughout this study, as well as the Koch Institute’s Robert A. Swanson (1969) Biotechnology Center for technical support, particularly the Barbara K. Ostrom (1978) Integrated Genomics and Bioinformatics. This study was supported by the National Institutes of Health (NCI) Koch Institute Support (core) grant P30-CA014051 and awards to: CEF (NIH NIGMS training grant award 5T32GM136540, and David H. Koch Graduate Fellowship), PLB (NIH GM141544) and JAL (NIH NCI-2P01CA04206331).

## Author contributions

C.E.F. and J.A.L. conceived this study, with input from PLB, and wrote the manuscript.

C.E.F conducted all of the experiments with assistance from: N.A.O. for the phenotypic characterization of cell lines and G.J.S. for RNA-Seq sample preparation and analysis. P.L.B. generated the computational pipeline with assistance from A.S.

## STAR Methods

### Cell Lines, Cell Line Generation and Inhibitors

U87, H23, PANC1, CAL51, HCT116, HEK293T, and RPE-1 cells were obtained from ATCC. Murine KP LUAD lines KP1233 and KP393T5 cell lines are previously described^40,41^. Murine KP PDAC line MDM1402 is previously described^42^. All cells were cultured in Dulbecco’s modified Eagle’s media (DMEM) supplemented with 10% fetal bovine serum (FBS) and 1% penicillin-streptomycin, and cultured in 37℃ at 5% CO_2_. Η23 cells were cultured in RPMI-1640 media supplemented with 10% fetal bovine serum (FBS) and 1% penicillin-streptomycin, and cultured in 37℃ at 5% CO_2_. Cells were routinely tested for mycoplasma and were found to be negative. The PRMT5 inhibitor JNJ-64619178 was obtained from MedChemExpress (HY- 101564), and the PRMT5 inhibitor EPZ015666 was obtained from L. Garraway. The dTag inducer molecule dTag-13 was obtained from Tocris (6605).

#### dTag-CLNS1A Knock-In Vector Generation

dTag (pCRIS-PITChv2-Puro-dTAG and pCRIS-PITChv2-BSD-dTAG) and PITCh (pX330A-1×2 and pX330S-2-PITCh) vectors were obtained from Addgene (91792, 91793, 58766, 63670). Both Puro and BSD dTag plasmids were modified to harbor homology to the N-terminus of CLNS1A as previously described^26^, using the following primers for Gibson Assembly:

dTag-CLNS1A Gibson F:

GTTCCGCGTTACATAGCATCGTACGCGTACGTGTTTGGTGGAGCGATGCTTTCCCCGGCTATGGCCAAGCCTTTGTCTCAAG

dTag-CLNS1A Gibson R:

AGCATTCTAGAGCATCGTACGCGTACGTGTTTGGGCTTTTGAGGAAGCTCATTGCTGGACTAAGCATAGATCCGCCGCCACC

The PITCh vector (pX330A-1×2) was modified to contain an sgRNA targeting the N-terminus of CLNS1A as previously described^43^, using the following primers:

sgCLNS1A NT F: CACCGTTTGAGGAAGCTCATTGCCG

sgCLNS1A NT R: AAACCGGCAATGAGCTTCCTCAAAC

Finally, pX330A-CLNS1A/PITCh was generated using Golden Gate Assembly.

#### dTag-CLNS1A Knock-In Cell Line Generation

Knock-in cell lines were generated as previously described^26^. Briefly, 10^6^ cells were plated in a 10-cm dish prior to transfection. The following day, cells were transfected with 2 μg pX330A- CLNS1A/PITCh and 1 μg pCRIS-PITChv2-Puro-dTAG-CLNS1A using Lipofectamine 2000 (ThermoFisher, 11668027). After 8 hours, media was replaced and supplemented with 500 nM NU7441 (Selleck Chemicals, S2638) to inhibit NHEJ. CAL51 cells did not tolerate NU7441 treatment, so this addition was omitted for that cell line. After 48 hours, cells were selected with puromycin. Following selection and outgrowth, cells underwent a second transfection with 2 μg pX330A-CLNS1A/PITCh and 1 μg pCRIS-PITChv2-BSD-dTAG-CLNS1A, using the same transfection workflow to enrich for homozygous knock-ins. Following blasticidin selection, cells were single cell cloned by dilution. CLNS1A knock-in was confirmed by gDNA PCR and western blot analysis.

#### Knockout Cell Line Generation

LentiCRISPRv2 and lentiCRISPRv2 Hygro were obtained from Addgene (98290 and 98291). Vectors were modified to contain an sgRNA targeting MTAP or CLNS1A using the following primers:

sgMTAP-1 F: CACCGTCTGCCCGGGAGCTAAAACG

sgMTAP-1 R: AAACCGTTTTAGCTCCCGGGCAGAC

sgMTAP-2 F: CACCGAAATACCATACCTTGCAAGG

sgMTAP-2 R: AAACCCTTGCAAGGTATGGTATTTC

sgCLNS1A-1 F: CACCGGAGGATTCAGATGACTACGA

sgCLNS1A-1 R: AAACTCGTAGTCATCTGAATCCTCC

sgCLNS1A-2 F: CACCGTCACACGCTGATTTATCACT

sgCLNS1A-2 R: AAACAGTGATAAATCAGCGTGTGAC

sgCtrl F: CACCGGTATTACTGATATTGGTGGG

sgCtrl R: AAACCCCACCAATATCAGTAATACC

For cell line generation, cells were infected with lentivirus containing lentiCRISPRv2 or lentiCRISPRv2 Hygro with a sgRNA targeting the gene of interest, or a control sgRNA (sgCtrl). Virus was incubated with cells for 2 days, after which puromycin or hygromycin was added for selection. Following selection cells were single cell cloned by dilution. For MTAP cell lines, knockout was confirmed by western blot analysis. For CLNS1A knockout cell lines, the CLNS1A locus was amplified and subcloned by Gibson Assembly into pcDNA3.1+ (ThermoFisher, V79020):

sgCLNS1A-1 Gibson F: TTGGTACCGAGCTCGTTCACTAACCATGTGCTTCCA

sgCLNS1A-1 Gibson R: GCTGGATATCTGCATGGCGTTTTGTGTACCCATTAAA

sgCLNS1A-2 Gibson F: TTGGTACCGAGCTCGTGGAGGTAATGGTGGTAAAGGTG

sgCLNS1A-2 Gibson R: GCTGGATATCTGCAAATTCATTCCCGGTGAGGCG

Following transformation into DH5α cells, colonies were sequenced by Sanger Sequencing. CLNS1A knockout was additionally confirmed by western blot analysis.

#### Overexpression Cell Line Generation

Overexpression studies constructs utilized the pUltra system, which we modified to incorporate a blasticidin resistance cassette. Briefly, pUltra Hot was digested with XbaI and AgeI-HF, and DNA containing the antibiotic resistance cassette introduced via Gibson assembly, generating the pUltra Blast vector. Full length human CLNS1A was ordered as a gBlock (IDT). CLNS1A Truncation was cloned from cDNA. Genes were cloned into the pUltra backbone via Gibson Assembly, which also incorporated a 3xFLAG-tag at the C-terminus of the gene.

Cells were infected with lentivirus to overexpress the gene of interest, or an empty vector control. Virus was incubated with cells for 2 days, after which antibiotic was added to generate stable cell lines.

### RNA Extraction, Sequencing and Alternative Splicing Analysis

#### Whole Cell Lysate RNA Extraction

Following 3-day treatment with either DMSO (vehicle), 10 nM PRMT5i, or 1 μM dTag-13, cells were harvested by scraping. RNA was purified using the RNeasy Mini Kit (Qiagen, 74106) according to manufacturer’s instructions, including the optional DNaseI addition to remove contaminating gDNA (Qiagen, 79254).

#### Cellular Fractionation and RNA Isolation from Cellular Fractions

U87 cells were seeded in a 15 cm plate at a 1.5×10^6^ cell/plate. One day after plating, cells were treated with vehicle (DMSO) or 10 nM PRMT5i (JNJ-64619178). After 3 days in drug, cells were harvested via trypsinization, pelleted, and separated into nuclear and cytoplasmic fractions as previously described^44^. Briefly, cell pellets were resuspended in cytoplasmic lysis buffer [0.15% (v/v) NP40, 10 mM Tris (pH7), 150 mM NaCl] supplemented with 10 U SUPERase.IN (Thermo Fisher, AM2694) and EDTA-free protease inhibitor cocktail (MilliporeSigma, 11836170001). Cells were then incubated on ice for 5 minutes. Cell lysate was layered on top of sucrose buffer [0 mM Tris (pH7), 150 mM NaCl, 25% (w/v) sucrose] supplemented with 10 U SUPERase.IN (Thermo Fisher, AM2694) and EDTA-free protease inhibitor cocktail (MilliporeSigma, 11836170001). Cells were then centrifuged at 16,000 x g for 10 minutes at 4 °C. The supernatant contains the cytoplasmic fraction and was harvested for RNA extraction. The remaining pellet contains the nuclear fraction and was washed twice with 800 μL cold PBS, centrifuging at 16,000 x g for 1 minute at 4°C after each wash. The pellet was resuspended in cold RIPA buffer [50 mM Tris (pH 8), 150 mM NaCl, 1% (v/v) NP-40, 0.5% sodium deoxycholate, and 0.1% (w/v) SDS] supplemented with 10 U SUPERase.IN (ThermoFisher, AM2694) and EDTA-free protease inhibitor cocktail (MilliporeSigma, 11836170001). Lysate was incubated for 30 minutes on ice, with vortexing every 10 minutes, and then pelleted at 16,000 x g for 10 minutes at 4°C. The supernatant contains the nuclear fraction and was harvested for RNA extraction.

The RNA from the nuclear and cytoplasmic fractions were extracted using TRIzol Reagent (Thermo Fisher, 15596026) according to manufacturer’s instructions. Briefly, to each aqueous fraction, 0.5 mL of isopropanol and 1 mL of TRIzol Reagent was added, followed by an incubation on ice for 10 minutes. Samples were then centrifuged for 10 minutes at 12,000 x g at 4 °C, pellets were washed in 1 mL 75% ethanol, and centrifuged again for 5 minutes at 7,500xg at 4 °C. Pellets were air dried for 10 minutes and resuspended in RNase-free water.

#### Library Preparation and Sequencing

For the human cell lines, RNA-Seq libraries were prepared using the NEB Ultra II Directional RNA Kit with Poly(A) Selection and sequenced using the NovaSeq sequencer, with 150 nucleotide paired-end reads. For the murine LUAD KP1233 cells, RNA-Seq libraries were prepared using the NEB Ultra II Directional RNA Kit with Poly(A) Selection and sequenced using the Element AVITI sequencer, with 75 nucleotide paired-end reads. For the murine PDAC and LUAD 393T5 cells, RNA-Seq libraries were prepared using the EB Ultra II Directional RNA Kit with Poly(A) Selection and sequenced using the NextSeq500, with 75 nucleotide paired-end reads.

#### Alternative Splicing Analysis

Raw RNA-seq reads were mapped using STAR aligner version 2.5.3a^45^ with the following parameters:

STAR

--runMode alignReads

--runThreadN 16

--sjdbOverhang 149

--outReadsUnmapped Fastx

--outSAMtype BAM SortedByCoordinate

--outFilterMultimapNmax 20

--outFilterMismatchNmax 999

--outFilterMismatchNoverLmax 0.04

--alignIntronMin 70

--alignIntronMax 500000

--alignMatesGapMax 500000

--alignSJoverhangMin 8

--alignSJDBoverhangMin 1

--outSAMstrandField intronMotif

--outFilterType BySJout

--twopassMode Basic

Additionally, raw RNA-seq reads were mapped using Bowtie 1.2.3^46^ using the following parameters:

bowtie-n 1-l 30-e 70-t-p 4-m 1-X 500-S

Detained intron and alternative splicing event identification was performed as previously described^2,21^, with the following modifications: Mapped reads were filtered using Bedtools v2.26.0 to remove reads overlapping expressed repeats from the UCSC genome browser RNA repeats and Gencode hg38 or mm10 and coding exons annotated in the Gencode hg38 or mm10 annotations. Alternative and constitutive classifications were performed using custom Python scripts and are agnostic with regard to existing annotations other than known gene boundaries. Importantly, all cell lines were run in parallel to ensure that the annotations would be consistent between the cell lines.

#### Detained Intron and Alternative Splicing Event Quantification (DEXSeq)

Detained intron and alternative splicing events were quantified using DEXSeq^47^. A genome annotation including all alternative splicing events and detained introns was generated using custom python scripts and python scripts that accompany DEXSeq. Reads were then quantified to this genome using the DEXSeq python script with the following parameters:

-p yes-s reverse-f bam-r pos

Finally, differential splicing was quantified using the DEXSeq package within R. Alternative splicing events and detained introns were called as significant if p_adj_<0.05 and lowly expressed exons and introns were filtered out if their normalized counts were below 80.

#### Detained Intron and Coding Isoform Quantification (RSEM-DESeq2)

Transcript annotation was performed as previously described^2^. Briefly, using custom Python scripts, we derived a complete transcriptome annotation based on Gencode hg38 gene start and end boundaries and the location of polyA sites for data derived from the human cell lines, the genomic locations of detained introns (as determined above), and the genomic locations of all splice junctions (from the STAR alignment). For comparisons of isoforms containing detained introns to the consensus coding sequence, the annotation for each gene was set to identify the consensus isoform, i.e., the junctions defining each exon are the most frequently used junction detected within all of the combined samples. Each detained intron within a gene was then assigned to an additional transcript.

#### Gene and Isoform Expression

Isoform-level quantification was performed using RSEM v1.3.1^48^ using the custom genomic annotation described above, as well as Gencode hg38 annotations with the parameters:

`rsem-calculate-expression--forward-prob 0--no-bam-output --paired-end` Differential expression analysis of gene and isoform count tables was performed using DESeq2^49^. Normalized count tables were obtained from DESeq2. Gene and isoform expression differences were considered significant if p_adj_<0.01.

#### Dice Similarity Score

Dice Similarity scores were calculated for each AS event type across all pairwise cell line combinations using the following equation:

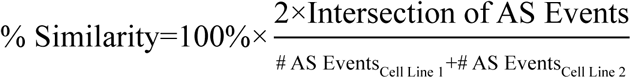

#### Gene Ontology Analysis

Gene Ontology (GO) analysis was conducted using DAVID^50,51^. Genes who contained a significant DI upon PRMT5i (p_adj_<0.05, log_2_FC>0) were run against the KW Biological Process category. Gene sets were considered significant if FDR<0.01. Gene sets are displayed if the med the significance threshold in at least one line.

#### Inter-species Gene Name Conversion and Complex Analysis

Lists of genes containing a significant DI upon PRMT5i (p_adj_<0.05, log_2_FC>0) for each cell line of interest was created. Mouse gene annotations were converted to human gene annotations using NicheNet v2.1.7^52^. The Comprehensive Resource of Mammalian Protein Complexes (CORUM) v4.1^38^ was used to match DI-containing genes to annotated human protein complex members. Gene Ontology (GO) analysis was conducted using DAVID as described above. The DI-containing genes for each protein complex were used to connect complexes to related biological processes. We directly compared PRMT5 DI-regulated genes, protein complexes, and biological processes between human and mouse data based on tissue of origin (Lung: H23 & T5, Pancreas: PANC1 & PDAC), as well as data derived from PRMT5 DI-containing genes shared between all human cell lines (n=7) or all mouse cell lines (n=3). Similarities were determined via a Dice Similarity Score as described above.

### siRNA Knockdown

U87 cells were seeded cells in a 6 well dish at a 10^5^ cells/well. One day after plating, cells were treated with vehicle (DMSO) or 10 nM PRMT5i (JNJ-64619178). After one day in drug, cells were transfected with siRNAs targeting UPF1 (Sigma-Aldrich, SASI_Hs01_00101017 and SASI_Hs01_00101018) or a non-targeting siRNA control (Sigma-Aldrich, SIC001) at a 10 nM concentration with 12 μL of MISSION siRNA Transfection Reagent (Sigma-Aldrich, S1452) for a final volume of 2.2 mL per well. Cells were harvested via trypsinization 48 hours post transfection and RNA was extracted as described above.

### Western Blot

Cells were harvested by scraping, washed twice with PBS, and then lysed in RIPA buffer [50 mM Tris pH 8.0, 150 mM NaCl, 1% NP-40, 0.5% sodium deoxycholate, and 0.1% sodium dodecyl sulfate (SDS)] supplemented with EDTA-free protease inhibitor cocktail (MilliporeSigma, 11836170001). For drug treatment experiments, cells were treated with DMSO, PRMT5i, or dTag-13 at the indicated concentration for the indicated duration, before harvesting. Protein concentration was determined using the Pierce BCA Protein Assay Kit (Thermo Fisher, 23227), then 30 μg of protein was mixed with 4x Laemmli buffer and loaded onto 12% SDS-polyacrylamide gels. Proteins were separated by SDS-PAGE, transferred onto nitrocellulose membranes, and probed overnight at 4℃ in 5% non-fat milk in TBST (unless otherwise stated) with antibodies against the following: CLNS1A (Invitrogen, PA5-109555, 1:1000), HA (Cell Signaling Technology, 3724S, 1:5,000), GAPDH (Thermo Fisher, AM4300, 1:5000), SDMA (Cell Signaling Technology, 13222S, 1:1,000, 5% BSA), Hsp90 (BD, 610418, 1:1,000), MTAP (Cell Signaling Technology, 4158S, 1:1000), FLAG (Sigma, F3165, 1:2000). Membranes were then incubated for 1 hour at room temperature with anti-mouse (LI-COR Biosciences, 926-32210, 1:10,000) or anti-rabbit (LI-COR Biosciences, 926-32211, 1:10,000) conjugated to IRDye 800CW and fluorescence was measured using the ChemiDoc Imaging System (Bio-Rad). Blots were quantified using Image Lab 6.1 (Bio-Rad).

### RT-PCR and Quantitative Real-Time PCR (qRT-PCR)

Total RNA was purified from cultured cells using the Rneasy Mini Kit (Qiagen, 74106) according to manufacturer’s instructions, including the optional DNaseI addition (Qiagen, 79254). RNA was converted to cDNA using SuperScriptIII following the manufacturer’s instructions (Thermo Fisher, 18080044).

Quantitative Real-Time PCR (qRT-PCR) reactions were performed using FAST-SYBR Green (Thermo Fisher, 4385612) on a StepOnePlus Real-Time PCR System (Applied Biosystems). Data were analyzed using the ΔΔC_T_ method, and relative messenger RNA (mRNA) levels were normalized to *GAPDH*. The following primers were used:

*GAPDH* F: GACAGTCAGCCGCATCTTCT

*GAPDH* R: GCGCCCAATACGACCAAATC

*DNA2* DI F: GGATGGCTTCTTATTTCATGG

*DNA2* DI R: CAGGCGCTTTTCACAGTTTC

*EIF4E* DI F: TGCAAATACAGAAGAGACATTTGC

*EIF4E* DI R: ACAACAGCGCCACATACATC

*UPF1* F: CCTGGGCCTTAACAAGAA

*UPF1* R: CCGCATGTCTCTTAACCA

FL-*CLNS1A* F: CCTGTTGCTGATGAAGAAGAGGA

FL-*CLNS1A* R: ACATTGCCTCCAACGCTGAT

Truncated-*CLN1A* F: CCCCTGTCCTTGTTCTTCAAA

Truncated-*CLNS1A* R: GCCGCCTGTCTTGGTTAGAT

RT-PCR reactions were performed using Phusion using the following primers:

CLNS1A F: GGCTTGGCCTTCTGCTGTTA

CLNS1A R: GCCGCCTGTCTTGGTTAGAT

Following amplification, PCR products were run on a 2% agarose gel, which was imaged using the ChemiDoc Imaging System (Bio-Rad).

### Dose-Response Curves

For each dose response curve, cells were plated in a volume of 150 μL media into each well of a 96-well plate in the following densities: CAL51 – 250 cells/well, HEK293T – 100 cells/well, and HCT116 – 150 cells/well. Perimeter wells were filled with 200 μL of media to avoid edge effects. The following day, 50 μL of 4x the final drug concentration in media was added to each well in triplicate. After 6 days of treatment, CellTiter-Glo (Promega, G7570) was used per manufacturer’s instructions. Luminescence was measured with a Tecan M200 Pro plate reader. Background signal was determined by averaging the luminescence of cell-free wells, and was subtracted from all other viability measurements. Average signal for each dose triplicate was computed, and drug-treated cells were normalized to their DMSO control by division. To generate dose-response curves, normalized viability values for each sample were fit to the following nonlinear model:

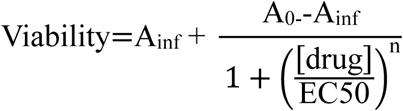

where A_inf_ and A_0_ are the lowest and highest drug doses, respectively. Area under the curve was computed and used for statistical testing.

### Proliferation Assay

For each cell line, 150 cells were plated in each well. Two replicate plates were plated. The following day, a plate was washed and frozen to serve as counting control. After 6 days, the other plate was washed and frozen. Cyquant was used according to the manufacturer’s instructions (ThermoFisher, C7026). Fluorescence was measured with a Tecan M200 Pro plate reader. Background signal was determined by averaging the fluorescence of cell-free wells and was subtracted from all other measurements. Population doublings were calculated via log-transformation of normalized fluorescence.

### Alphafold2 Prediction

Full length or Truncated CLNS1A protein sequences were inputted into the Alphafold2 ColabFold v1.5.5 software^53^, using the default settings. Structure predictions were visualized using PyMOL v2.5.2.

### Statistical Analysis and Plotting Software

R 4.3.2 and GraphPad PRISM 10.1.0 were used to perform statistical analyses. R 4.3.2, GraphPad PRISM 10.1.0, and Adobe Illustrator 28.2 were used for generating and plotting figures.

---

**Supplemental Figure 1.**
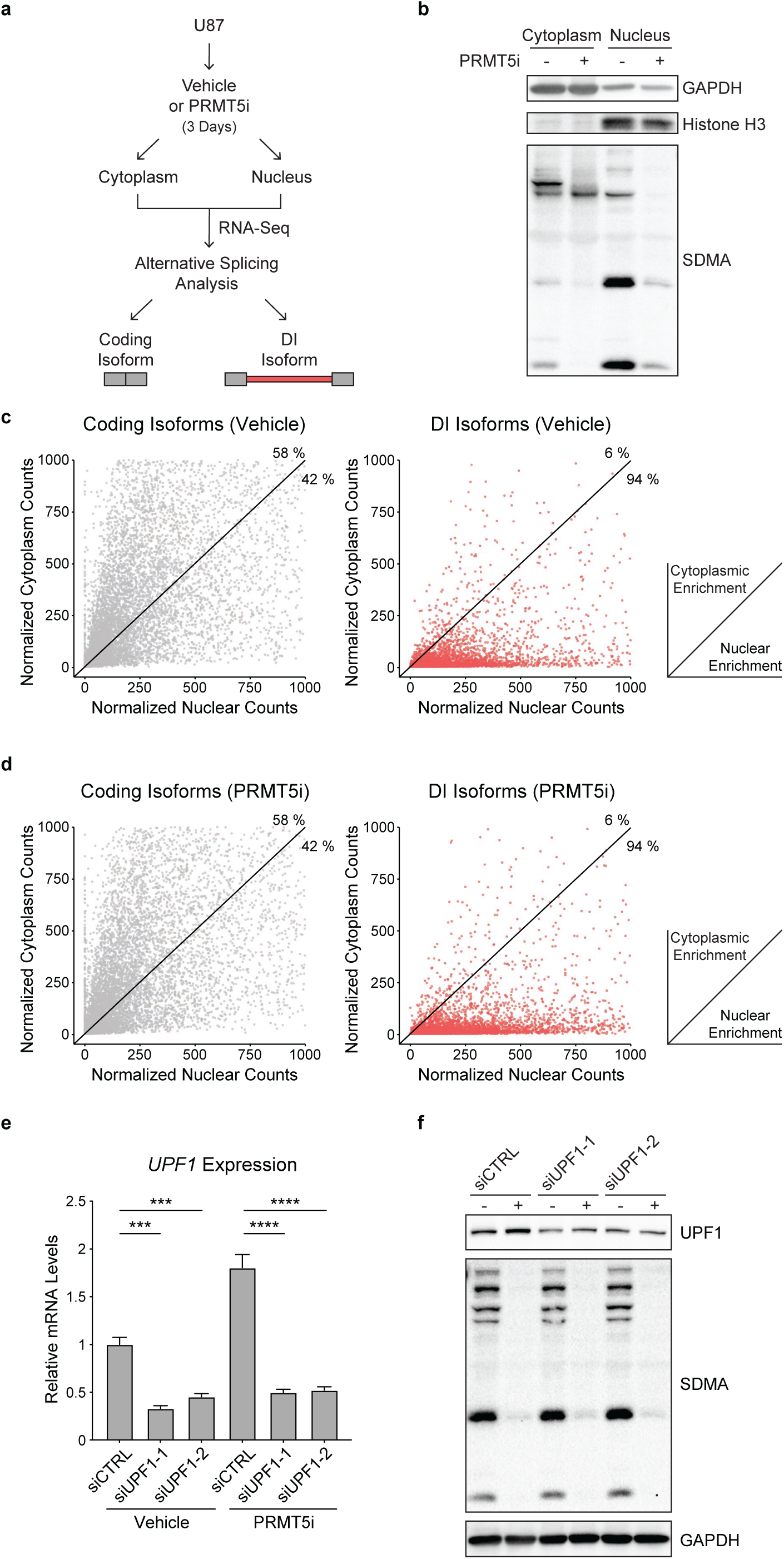
DI-containing transcripts show nuclear localization. **a.** Experimental schematic for the fractionation-coupled RNA-sequencing experiment. **b.** Western blot of nuclear and cytoplasmic fractions of U87 cells after 3-day treatment with vehicle control or 10 nM PRMT5i, using GAPDH and Histone H3 as cytoplasmic and nuclear markers, respectively. **c-d.** Quantification of coding (*Left*) versus DI-containing (*Right*) isoforms in cytoplasmic and nuclear fractions of U87 cells after 3-day (**c**) vehicle or (**d**) 10 nM PRMT5i treatment. Each point represents an individual isoform. Points above the line exhibit cytoplasmic enrichment, while ones below exhibit nuclear enrichment. Points along the axes are almost entirely cytoplasmic or nuclear. **e-f.** U87 cells were treated from 3-days with vehicle or 10 nM PRMT5i and transfected with two individual siRNAs targeting UPF1 or a non-targeting siRNA for the last 2 days prior to harvest. These samples were assessed for: **e.** Relative levels of UFP1 mRNA, as determined by qRT-PCR, (***p<0.001, ****p<0.0001, Student’s t-test); and **f.** levels of UPF1 protein, SDMA, and loading control GAPDH by western blotting.

**Supplemental Figure 2.**
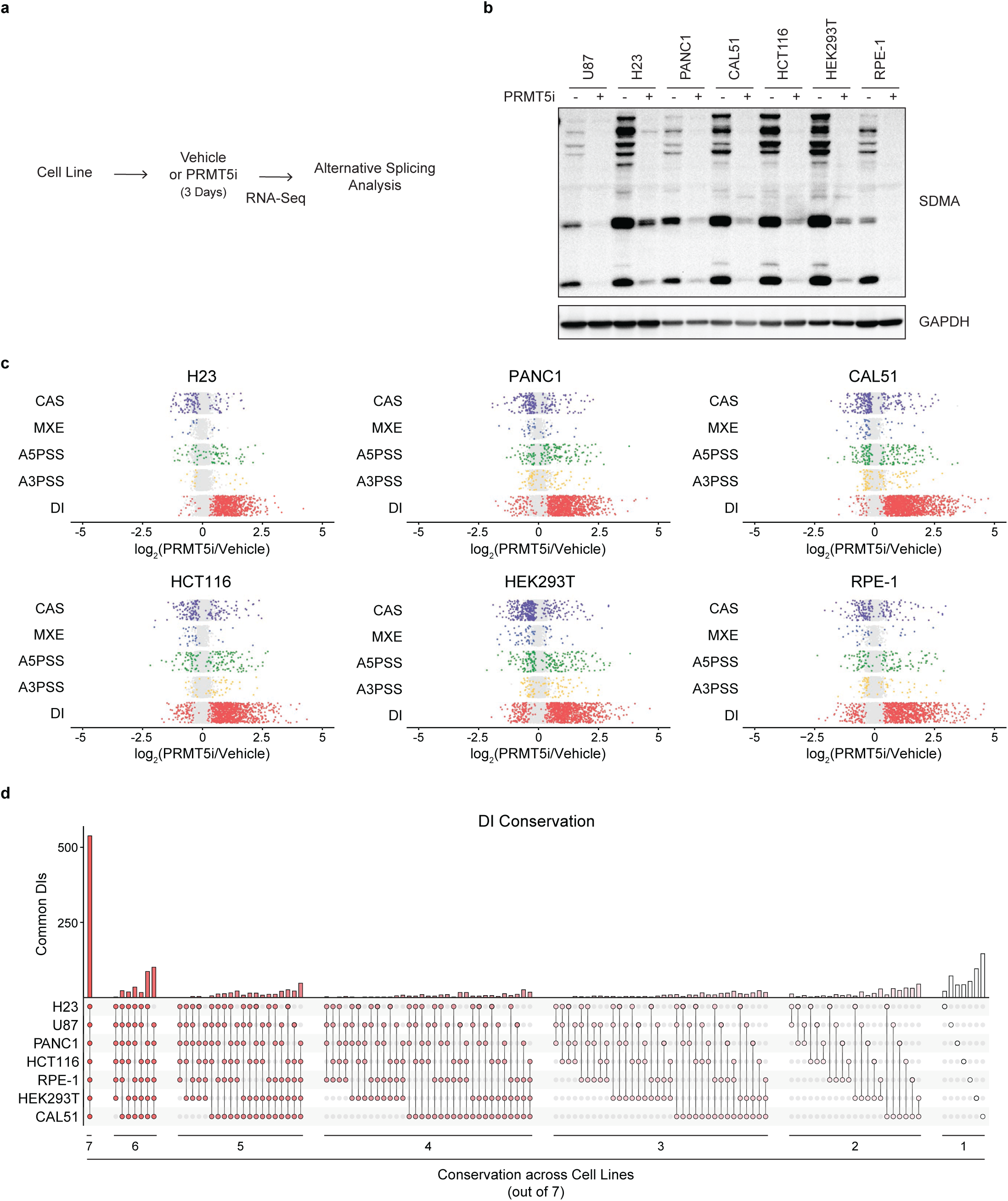
DIs are a conserved PRMT5-regulated AS event. **a.** Experimental schematic for the analysis of PRMT5i-responsive DIs in the human cell lines. **b.** Western blot of representative levels of SDMA and loading control GAPDH in indicated human cell lines with 3-day vehicle or 10 nM PRMT5i treatment. **c.** Induction of AS events in the indicated cell line after 3-day PRMT5i treatment compared to vehicle. Each row shows a specific AS event type with colored dots indicating significant events (p_adj_<0.05) and gray dots non-significant events. **d.** Upset plot of DI conservation across all human cell lines grouped by level of individual PRMT5- regulated DI conservation. Each column represents a unique conservation pattern where the colored dots below the x-axis indicate the cell lines with significant PRMT5-responsiveness. Bar height indicates the number of individual DIs that fit the conservation pattern.

**Supplemental Figure 3.**
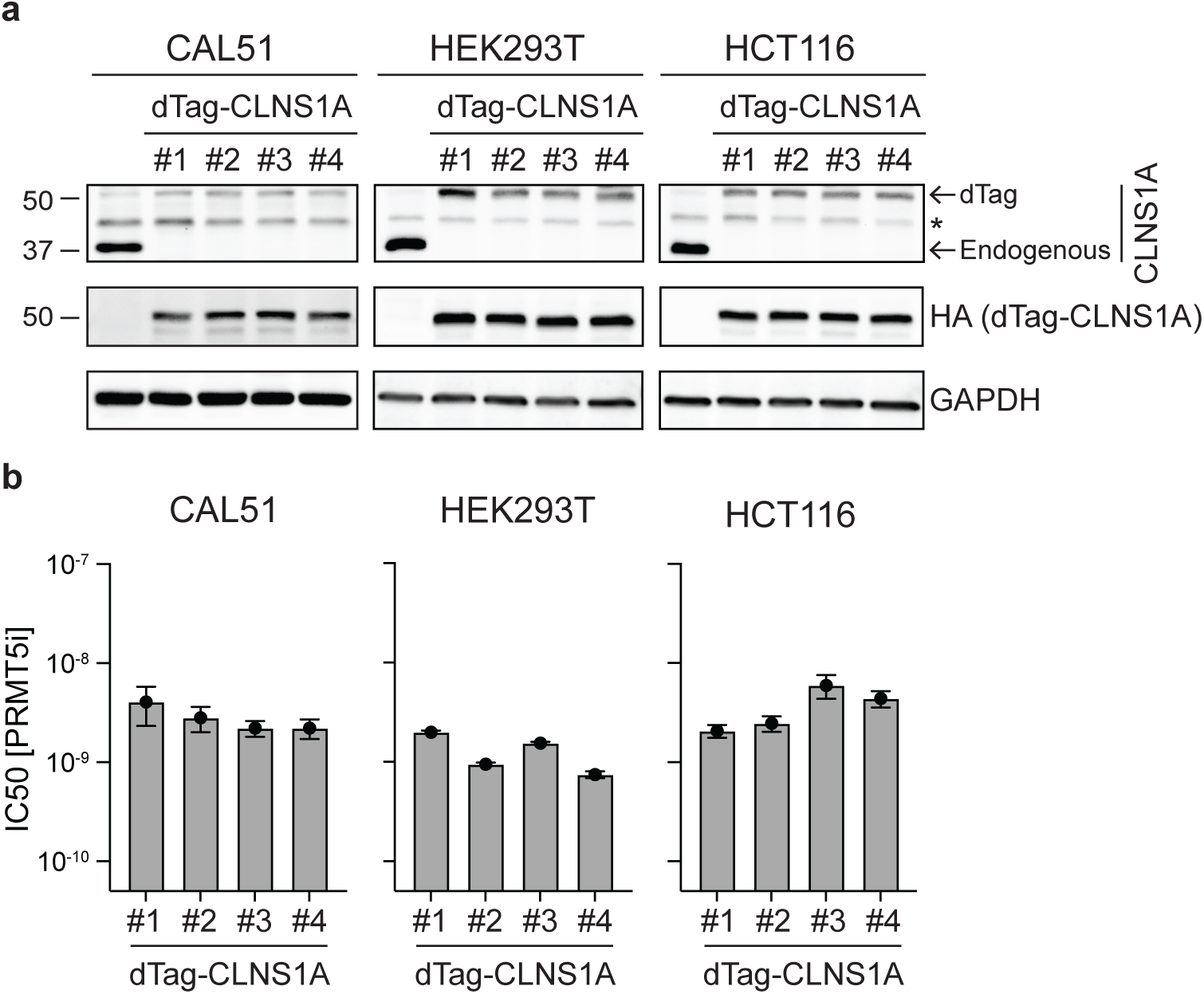
dTag-13 treatment selectively causes CLNS1A loss in engineered human cell lines. **a.** Western blot analysis of parental control and four dTag-CLNS1A clones for CAL51, HEK293T, and HCT116 cells with antibodies against CLNS1A, the inserted HA tag, and loading control GAPDH. Asterisk (*) marks a non-specific band. **b.** PRMT5i IC50 values for four representative dTag-CLNS1A clones derived from CAL51, HEK293T, and HCT116 cell lines. Data are mean ± SD of 3 technical replicates/line.

**Supplemental Figure 4.**
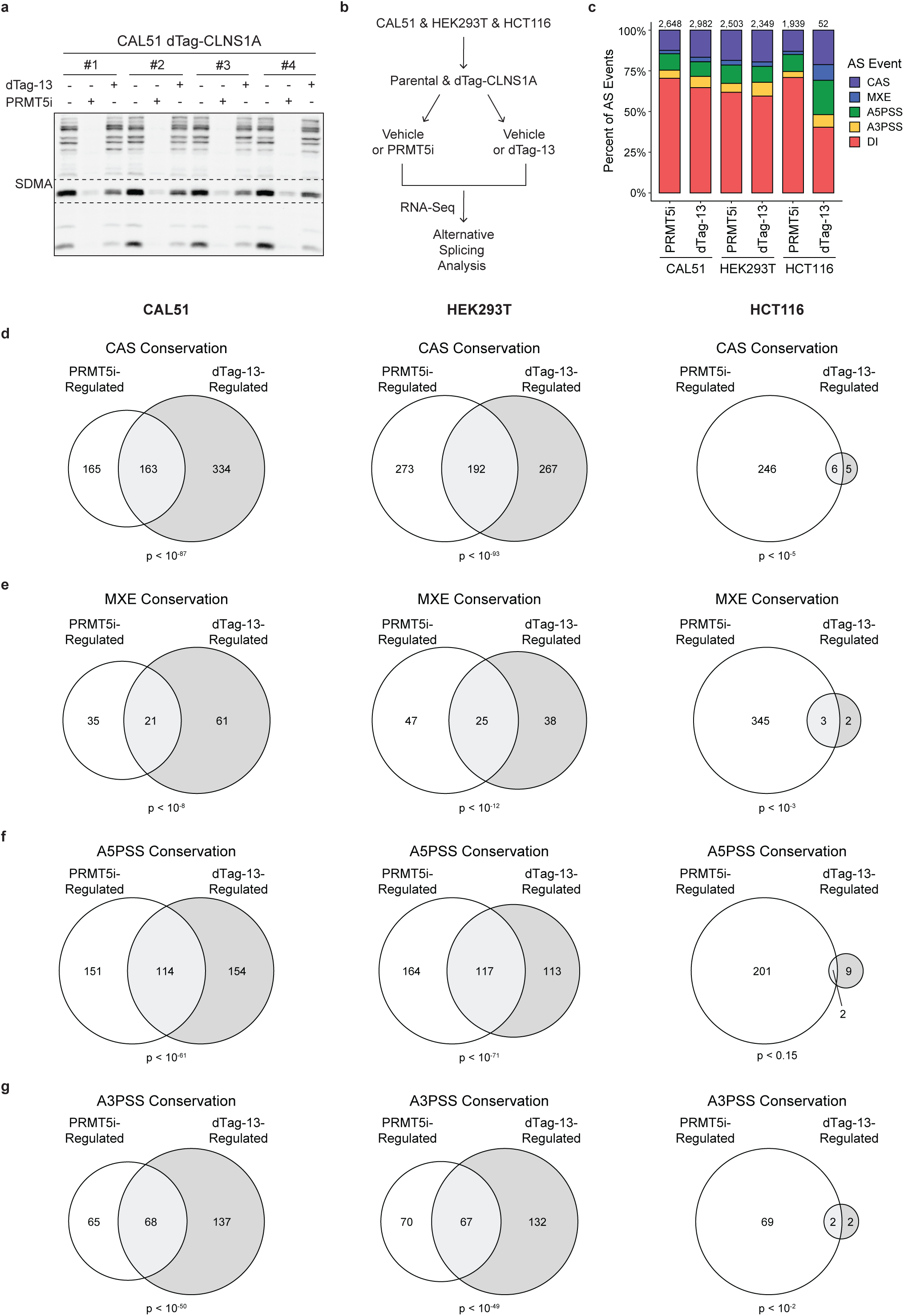
The PRMT5-splicing axis underlies AS events in dTag-13-sensitive lines. **a.** Uncropped western blot of the four CAL51 dTag-CLNS1A clones treated with either vehicle, 10 nM PRMT5i, or 1 μM dTag-13 for 3 days. Dashed lines signify where blot was cropped to incorporate into **Figure 4a**. **b.** Schematic of the dTag-13 and PRMT5i sequencing experiment used compare significant AS events between the two experimental conditions. **c.** Proportion of each AS event type that is significantly different in indicated cell line after 3-day treatment with 10 nM PRMT5i or 1 μΜ dTag-13 compared to vehicle control (p_adj_<0.05). Number above bar represents the total number of significant AS events. **d-g.** Overlap of (**d**) CAS, (**e**) MXE, **(f**) A3PSS, and (**g**) A5PSS events that were significantly altered by 3-day treatment with PRMT5i or dTag-13 compared to vehicle control (p_adj_<0.05) in the indicated CAL51 (*Left*), HEK293T (*Middle*), and HCT116 (*Right*) cells. Overlap significance was calculated using a hypergeometric test.

**Supplemental Figure 5.**
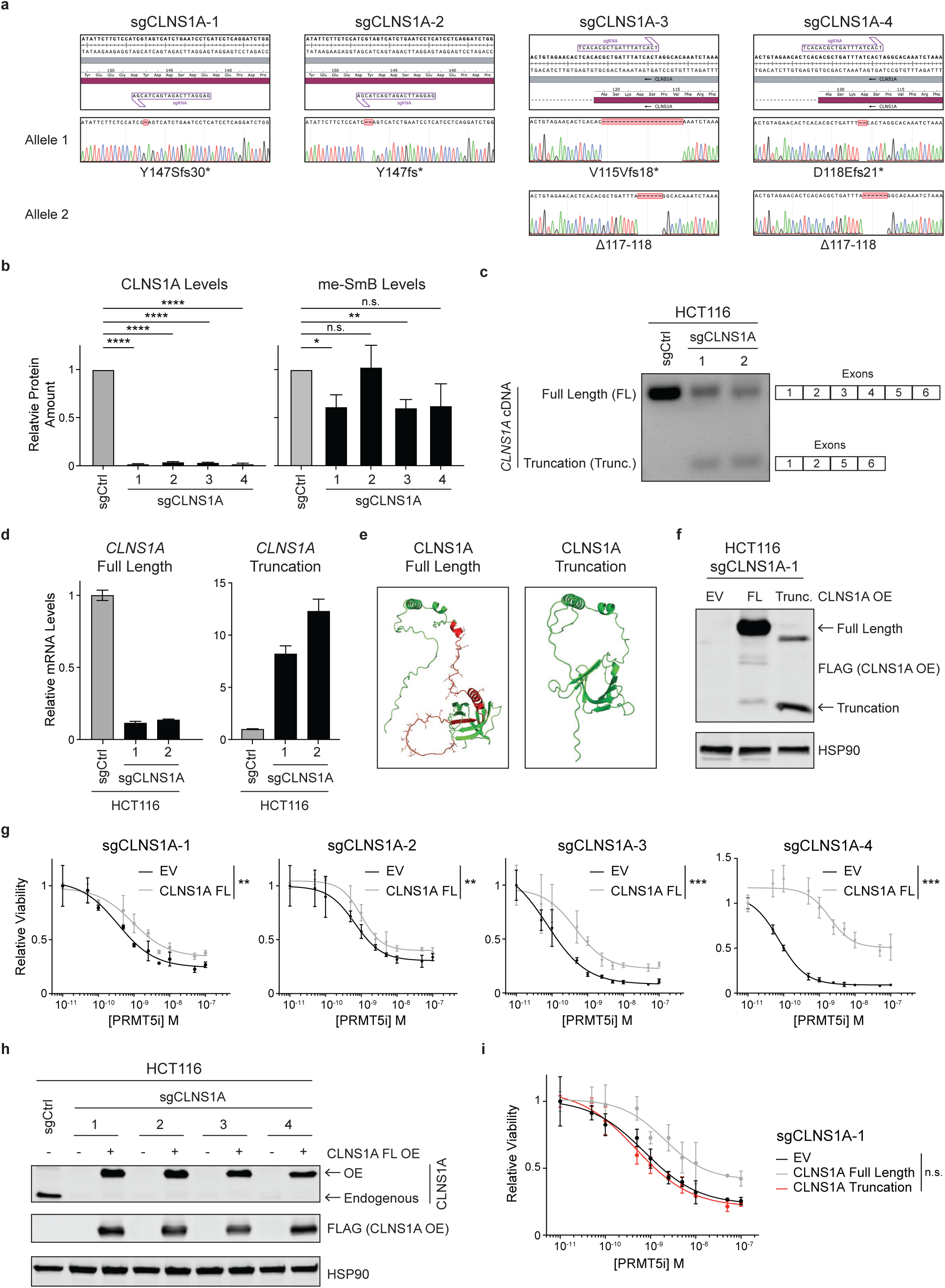
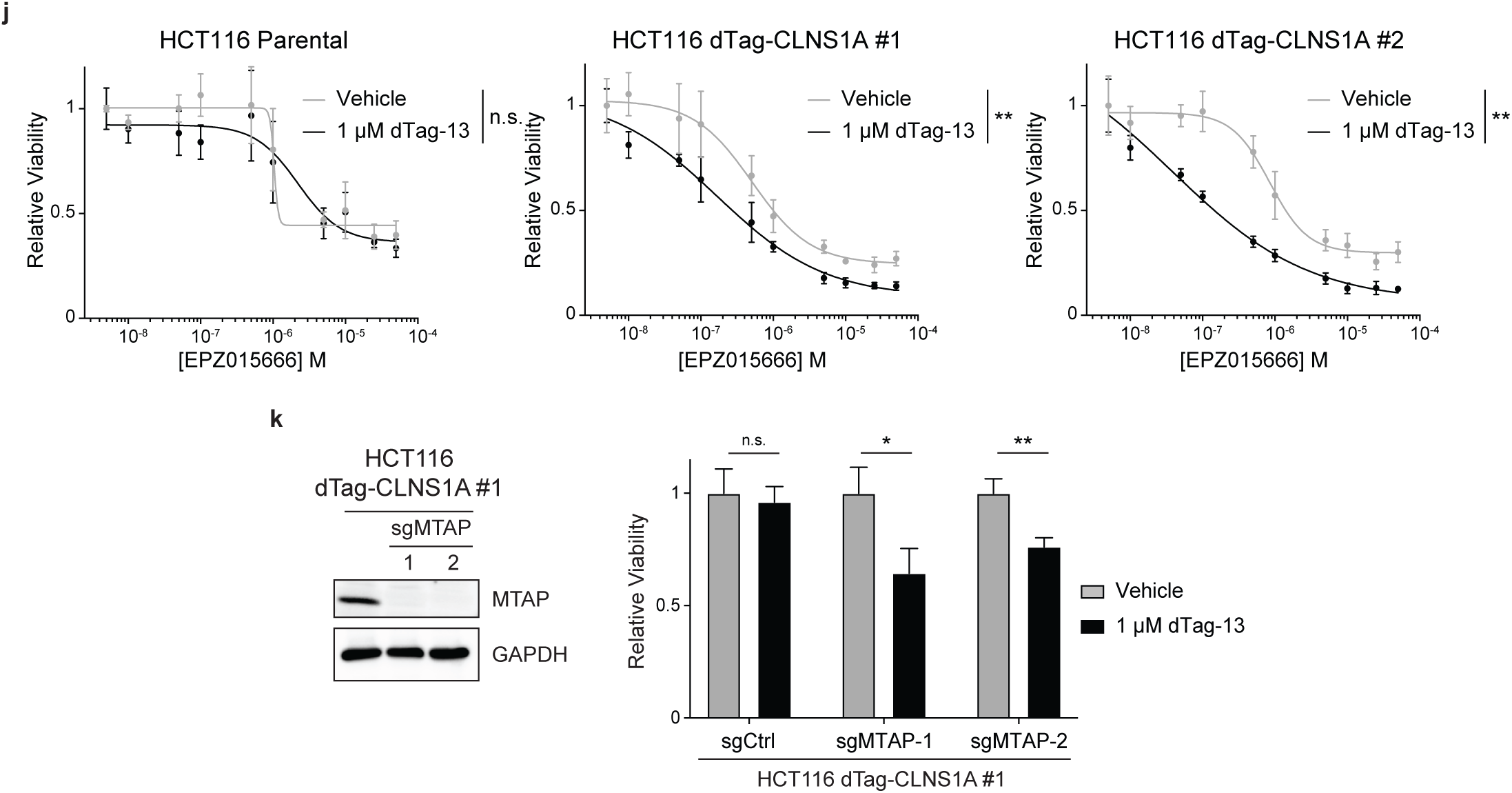
PRMT5 inhibition or MTAP deletion sensitizes cells to CLNS1A loss in otherwise CLNS1A-independent cells. **a.** Sanger sequencing traces of *CLNS1A* alleles in each of the four HCT116 sgCLNS1A clones. Expected protein product for each allele is indicated. **b.** Quantification of CLNS1A (*Left*) and methyl-SmB (*Right*) levels from **Figure 5b** and 3 other replicate blots, normalized to loading control. Data are mean ± SD of 4 biological replicates. *p<0.05, **p<0.01, ***p<0.001, and ****p<0.0001, Welch’s t-test. **c.** Agarose gel of RT-PCR products from HCT116 sgCtrl and sgCLNS1A-1 and-2 for the expression of FL- or truncated-*CLNS1A*. **d.** Relative mRNA expression of FL- and truncated-*CLNS1A* in HCT116 sgCtrl and sgCLNS1A- 1 and-2. Data represents mean ± SD of 3 technical replicates **e.** Alphafold2 models of FL- and truncated-CLNS1A. The red backbone indicates truncated amino acids, and acidic amino acids in this region are shown as sticks. **f.** Western blot analysis of FLAG-tag and loading control levels in HCT116 sgCLNS1A-1 infected by lentiviruses expressing empty vector (EV), FL-CLNS1A-3xFLAG, or truncated-CLNS1A- 3xFLAG. **g.** Dose response curve of HCT116 sgCLNS1A clones exogenously expressing EV control or FL- CLNS1A treated with JNJ-64619178 (PRMT5i) for 6 days. Data represents mean ± SD of 3 technical replicates. **p<0.01 and ***p<0.001, Student’s t-test. **h.** Western blot analysis of CLNS1A, FLAG-tag, and loading control levels in HCT116 sgCtrl and four sgCLNS1A clones exogenously expressing EV or FL-CLNS1A-3xFLAG. **i.** Dose response curve of the HCT116 sgCLNS1A-1 cell line carrying EV, FL-CLNS1A, or truncated-CLNS1A and treated with PRMT5i for 6 days. Data represents mean ± SD of 3 technical replicates. p=n.s., Student’s t-test. **j.** Dose response curve of the HCT116 parental or two dTag-CLNS1A clones treated with or without 1 µM dTag-13 in combination with the indicated concentrations of the PRMT5i EPZ015666 for 6 days. Data represents mean ± SD of 3 technical replicates. **p<0.01, Student’s t-test. **k.** Derivatives of HCT116 dTag-CLNS1A #1 cells expressing sgCtrl, sgMTAP-1 or sgMTAP-2 were analyzed for (*Left*) levels of MTAP and loading control GAPDH by western blot and (*Right*) relative viability after 6-day treatment with 1 μM dTag-13 or vehicle control. Viability data represents mean ± SD of 3 technical replicates. *p<0.05 and **p<0.01, Student’s t-test.

**Supplemental Figure 6.**
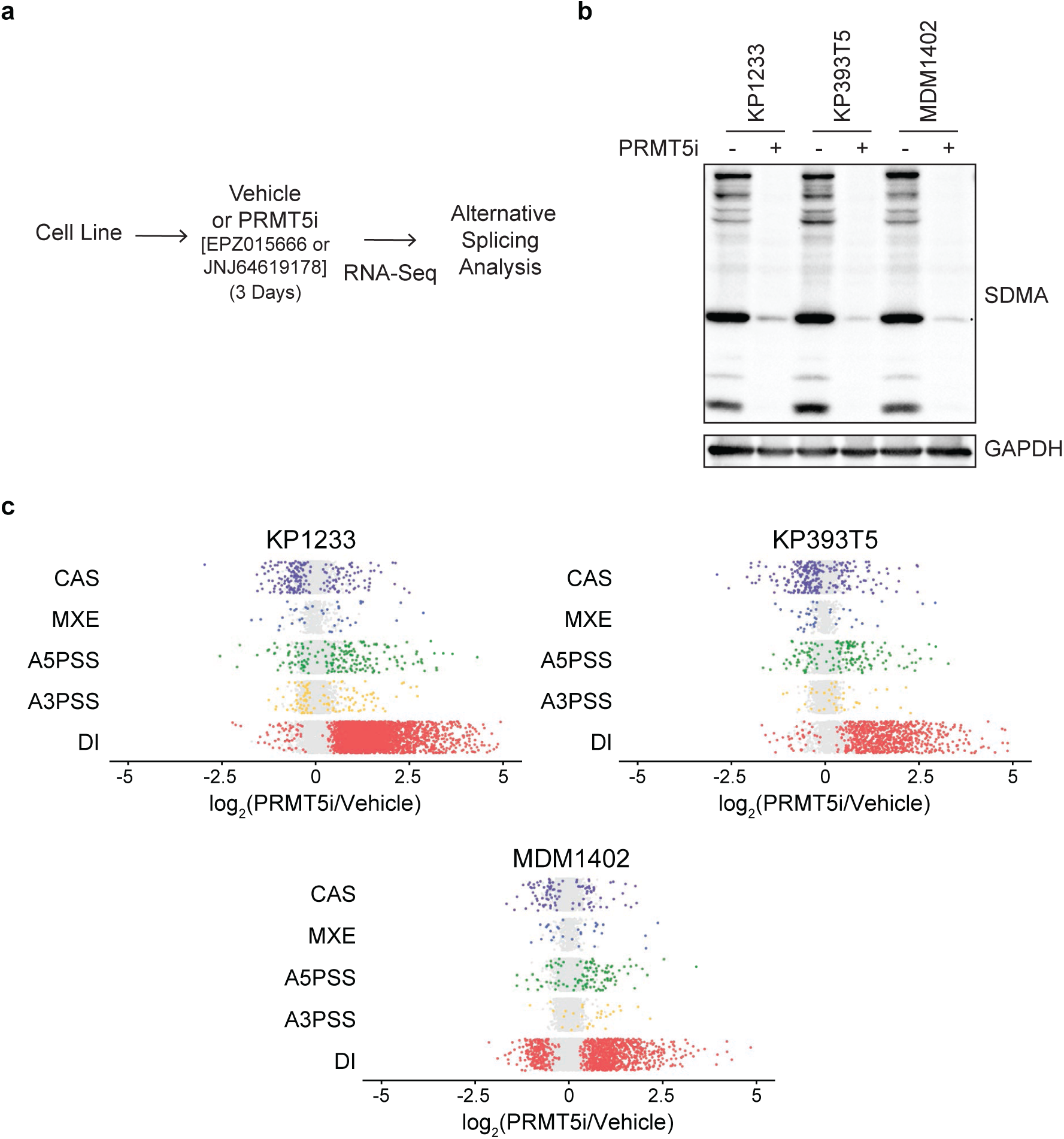
DIs are a conserved PRMT5-regulated AS event in mouse cancer cells. **a.** Experimental schematic for the analysis of PRMT5i-responsive DIs in murine cell lines. **b.** Western blot of representative SDMA levels and loading control GAPDH in indicated mouse cell lines after 3-day treatment with the PRMT5i indicated in **Figure 6a** or vehicle control. **c.** Induction of AS events in the indicated mouse cell line after 3-day PRMT5i treatment compared to vehicle. Each row is a specific AS event type. Colored dots represent significant events (p_adj_<0.05), while gray dots represent non-significant events.

**Supplemental Figure 7.**
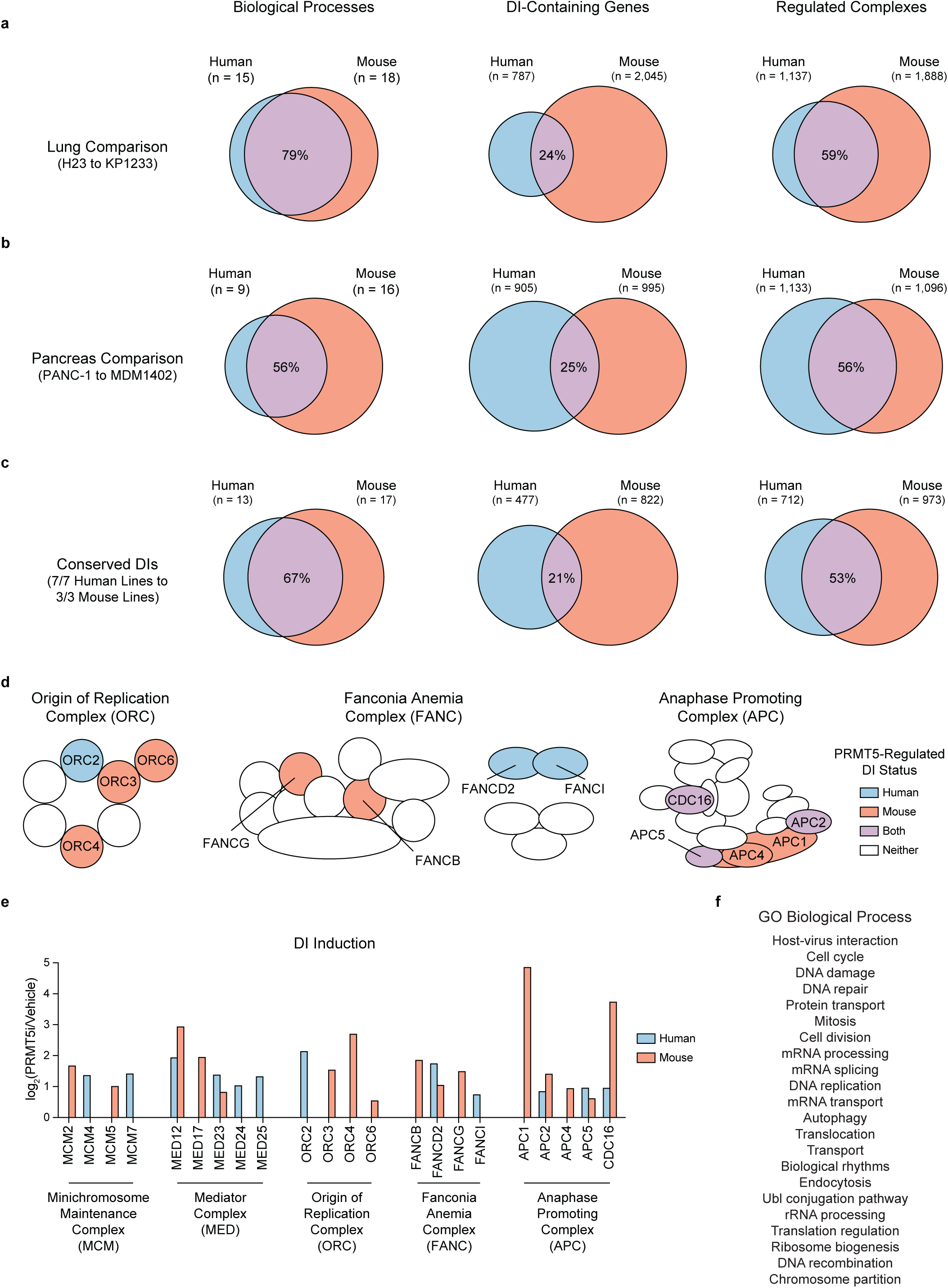
DI conservation increasingly converges to similar complexes and biological processes. **a-c.** Overlap of common (*Left*) DI-regulated processes, (*Center*) genes, and (*Right*) complexes for PRMT5-regulated DIs in (**a**) human (H23) and mouse (KP1233) lung cancer lines, (**b**) human (PANC1) and mouse (MDM1402) pancreatic cancer lines, and (**c**) all seven human or three mouse cell lines. Percentages represent the dice similarity score for each comparison. **d.** Schematic of representative DI-regulated complexes, including the origin of replication (ORC), Fanconi anemia (FANC), and anaphase promoting (APC) complexes. Each shape represents a specific complex member, and a color fill of blue, red, purple, or white indicates that the complex member contained a DI in human, mouse, both species, or neither, respectively. **e.** Induction of DIs in representative complexes shown in **Figure 7c** and **Figure S7d**. Each pair of columns represents a specific genes in the complex, where blue and red bars indicate the induction of DIs in human H23 and mouse KP393T5 lung cancer lines, respectively (p_adj_<0.05). Absence of a bar indicates no significant DI in either species. **f.** GO Biological Process categories for **Figure 7d** (*Column 3*) listed in order.

